# Synergistic control of chloroplast biogenesis by *MYB-related* and *Golden2-like* transcription factors

**DOI:** 10.1101/2023.08.11.552970

**Authors:** Eftychios Frangedakis, Nataliya E. Yelina, Kumari Billakurthi, Tina Schreier, Patrick J. Dickinson, Marta Tomaselli, Jim Haseloff, Julian M. Hibberd

## Abstract

Chloroplast biogenesis is dependent on master regulators from the GOLDEN2-LIKE (GLK) family of transcription factors, but *glk* mutants contain residual chlorophyll and therefore other proteins must also be involved. Here we identify MYB-related transcription factors as regulators of chloroplast biogenesis in the liverwort *Marchantia polymorpha* and angiosperm *Arabidopsis thaliana*. In both species, double mutant alleles in MYB-Related genes show very limited chloroplast development, and photosynthesis gene expression is perturbed to a greater extent than in mutants of GLK. In *M. polymorpha* MYB-related genes act upstream of GLK, while in *A. thaliana* this relationship has been rewired. In both species, genes encoding enzymes of chlorophyll biosynthesis are controlled by MYB-related and GLK proteins whilst those allowing CO_2_ fixation, photorespiration and photosystem assembly and repair require the MYB-related proteins. Thus, *MYB-related* and GLK genes have overlapping as well as distinct targets. We conclude that together MYB-related and GLK transcription factors orchestrate chloroplast development in land plants.

## Introduction

Photosynthesis is fundamental to life and in eukaryotes takes place in organelles known as chloroplasts. It is widely accepted that chloroplasts originated from endosymbiosis between a photosynthetic prokaryote and heterotrophic eukaryote that was initiated more than one billion years ago.^1–3^ Since then, significant elaborations to the control of photosynthesis gene expression have taken place. For example, in plants the majority of genes allowing chloroplast biogenesis are encoded in the nucleus such that thousands are post-translationally imported from cytosol to chloroplast.^4,5^ Despite these significant rearrangements to the genetics of photosynthesis in most plants including major crops, the photosynthetic process has not been optimised by natural selection.^6,7^ One current limitation to improving photosynthesis in crops is knowledge of the underlying gene regulatory networks.

The expression of photosynthesis associated nuclear genes is responsive to light and also to processes intrinsic to the cell. For example, in angiosperms light is required for chloroplast formation, but hormones amplify or repress this response.^8^ These exogenous and endogenous inputs are integrated by key transcriptional regulators belonging to the GOLDEN2-LIKE (GLK) and GATA families of transcription factors (GATA Nitrate-inducible Carbon metabolism-involved [GNC] and Cytokinin-Responsive GATA Factor 1 [CGA1].^8–12^ However, *glk* mutants in *Arabidopsis thaliana*^10^, rice^13^ and also non-seed plants such as *Physcomitrium patens*^14^ and *Marchantia polymorpha*^15^ still contain chlorophyll. Moreover, mutants lacking functional GLK and GATA genes are not albino.^9,16^ In summary, other actors must allow assembly of the photosynthetic apparatus in the absence of these known regulators.

We therefore sought to identify new transcription factors acting alongside the master regulator GLK. As forward genetics has failed to identify such proteins, we rationalised that genetic redundancy had hindered their identification and that analysis of a species with a more compact genome would circumvent this issue. *Marchantia polymorpha* possesses a streamlined genome with many transcription factors represented by either one or two copies and the dominant form of the lifecycle is haploid.^17^ Moreover, control of greening is streamlined with only one copy of GLK being present, and orthologs of *GATAs* implicated in chloroplast biogenesis in other land plants^18^ not being required.^15^ We hypothesised that homologous transcription factors in *A. thaliana* and *M. polymorpha* act alongside GLK and so their expression should respond to light during photomorphogenesis in both species. After re-examination of publicly available RNA sequencing data, gene editing of transcription factors and detailed phenotypic analysis we identify two RR-MYB transcription factors as regulators of chloroplast biogenesis and photosynthesis gene expression in *M. polymorpha* and *A. thaliana*. In contrast to the GLK proteins that regulate expression of genes allowing chlorophyll biosynthesis and function of photosystem I and II, the RR-MYBs have a broader set of targets that extends to genes allowing CO_2_ fixation, photorespiration, photosystem assembly and repair. We conclude that these proteins function as master regulators of chloroplast biogenesis and photosynthesis gene expression. The data have implications for understanding chloroplast biogenesis and photosynthesis as well as other processes taking place in plastids such as nitrogen and sulphur assimilation, the biosynthesis of amino acids, fatty acids and carotenoids.

## Results

### MpRR-MYB5 regulates chloroplast development synergistically with its paralog MpRR-MYB2

We interrogated publicly available gene expression data sampled during the transition from non-photosynthetic to photosynthetic growth in *M. polymorpha*^19^ as well as *A. thaliana.*^20^ This identified 108 and 144 transcription factors that were upregulated after exposure to light in *M. polymorpha* and *A. thaliana* respectively **(Table S1 and S2)**. We then selected orthologs upregulated in both datasets with an unknown or chlorophyll-related annotation that were represented by a multigene family in *A. thaliana* **(Figure 1A)**. Fourteen candidates from *M. polymorpha* were identified **(Figure 1B and Table S2)**. Two of these (MpGLK, MpGATA4) are homologs of known photosynthesis regulators in *A. thaliana*^10,21^ and MpGLK has a confirmed role in *M. polymorpha.*^15^ The remainder included a number of B-BOX (BBX) domain proteins known to interact with the master regulator of photomorphogenesis HY5^22^, a homeobox-leucine zipper (HD-ZIP) protein ATHB17 whose ortholog in *A. thaliana* regulates photosynthesis-associated nuclear genes in response to abiotic stress,^23^ a C2H2 type zinc finger transcription factor with unknown function in *A. thaliana,* and a *MYB-related* gene predicted to regulate photosynthesis gene expression.^24,25^

**Figure 1:**
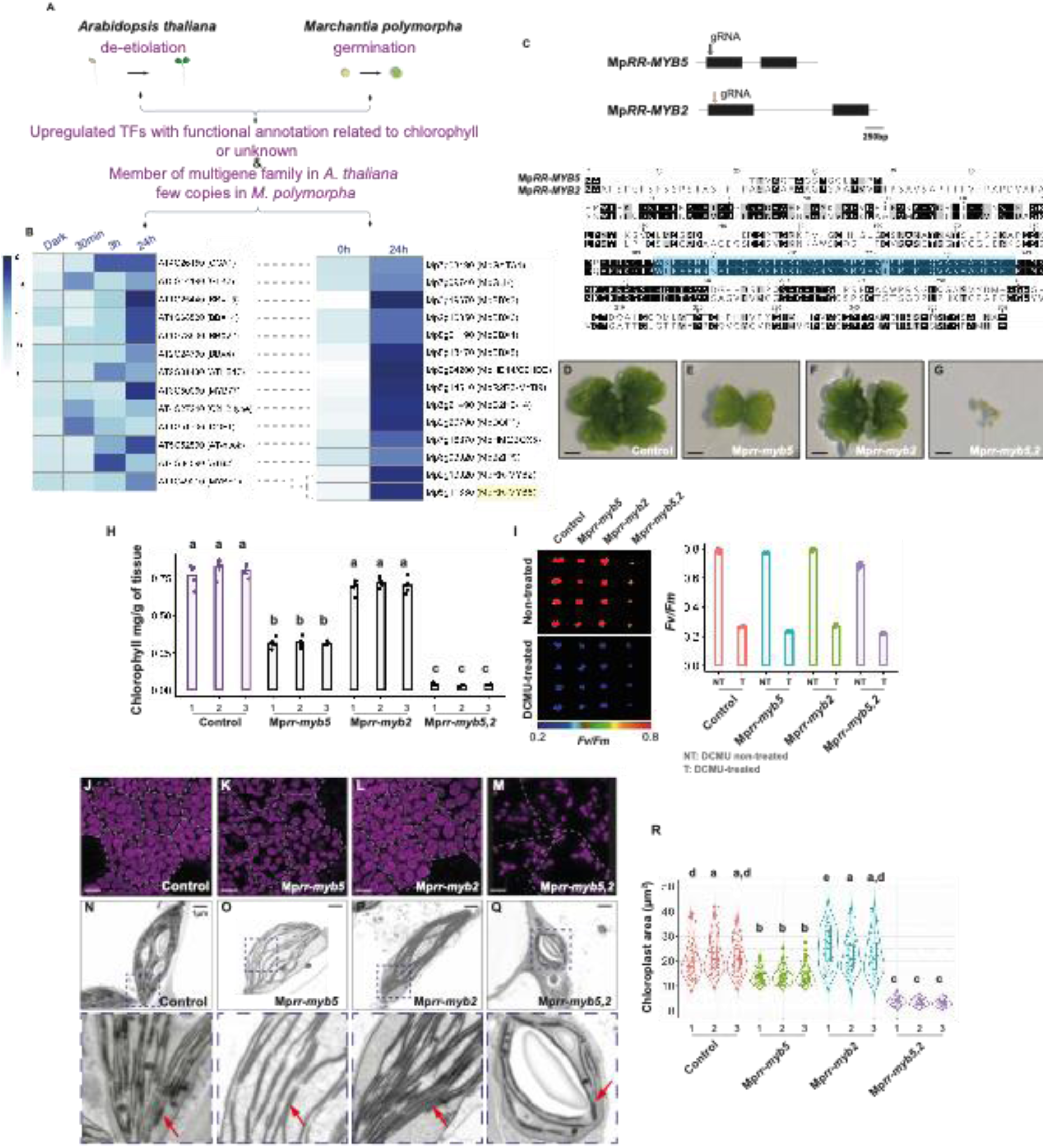
MpRR-MYB5 and MpRR-MYB2 act redundantly to regulate chloroplast biogenesis in *M. polymorpha*. **A)** Schematic of pipeline used to identify candidate transcription factors regulating photosynthesis gene expression. RNA sequencing data from *Arabidopsis thaliana* during de-etiolation^19^ and *Marchantia polymorpha* during spore germination^20^ were examined. Transcription factors (TFs) upregulated in response to light in both species with either unknown or related to chlorophyll function were retained. As we hypothesised that functional redundancy had hindered identification of such regulators via forward genetic screens an additional criterion was that each family should be represented by multiple copies in *A. thaliana* but by a maximum of two in *M. polymorpha*. **B)** Heatmaps showing transcript abundance (z-score) of candidates that were selected to generate knockout mutants in *M. polymorp*ha. **C)** Left: Schematic of MpRR-MYB5 and MpRR-MYB2 gene structure (exons represented as black boxes) with guide (g) RNAs for CRISPR represented by arrows. Right: Amino acid sequence alignments of MpRR-MYB5 and MpRR-MYB2 proteins with the characteristic RR-MYB/CCA1-like domain is highlighted in blue. **D-G)** Representative images of control and Mp*rr-myb5,* Mp*rr-myb2* and Mp*rr-myb5,2* mutants. Scale bars represent 2 mm. **H)** Chlorophyll content of Mp*rr-myb5*, Mp*rr-myb2* and Mp*rr-myb5,2* mutants. Letters show statistical ranking using a *post hoc* Tukey test with different letters indicating statistically significant differences at p<0.01. Values indicated by the same letter are not statistically different. n=5. **I)** Representative images and quantification after imaging of the chlorophyll fluorescence parameter *F_v_/F_m_* with and without the inhibitor Di-Chlorophenyl Di-methyl Urea (DCMU). Although the Mp*rr-myb5,* Mp*rr-myb2* and Mp*rr-myb5,2* mutants have lower chlorophyll content, the drop in *F_v_/F_m_* values after treatment with DCMU indicates that photosystem II is functional. **J-M)** Representative images after confocal laser scanning microscopy of Mp*rr-myb5*, Mp*rr-myb2* and Mp*rr-myb5,2* mutants with chlorophyll autofluorescence shown in magenta. Cell borders are marked with dashed white lines. Scale bars represent 10 μm. **N-Q)** Representative images after transmission electron microscopy of Mp*rr-myb5*, Mp*rr-myb2* and Mp*rr-myb5,2* mutants. Scale bars represent 1 μm. The dashed area depicted in each chloroplast is enlarged and grana stacks indicated with red arrows. **R)** Violin plots of chloroplast area for the Mp*rr-myb5,* Mp*rr-myb2* and Mp*rr-myb5,2* mutants. Box and whiskers represent the 25 to 75 percentile and minimum-maximum distributions of the data. Letters show statistical ranking using a *post hoc* Tukey test with different letters indicating statistically significant differences at p<0.01. Values indicated by the same letter are not statistically different, n=150.

Taking advantage of the rapid and predominantly haploid life-cycle we used *M. polymorpha* as a testbed for each of these candidates and subjected each to CRISPR/Cas9-mediated editing. With the exception of MpGLK, which has previously been reported to lead to a pale phenotype when mutated^15^ only one other candidate had low chlorophyll. This was Mp5g11830, annotated as MpRR-MYB5 in the *M. polymorpha* genome database, but previously also referred to as a CIRCADIAN CLOCK ASSOCIATED1-like RR-MYB-Related transcription factor.^26,27^ MpRR-MYB5 has a single paralog (MpRR-MYB2 - **Figure 1C**) that shows high similarity to MpRR-MYB5 at the amino acid level (**Figure 1C**) with for example the CCA1-like/RR-Myb domain being 92% identical (**Figure 1C**). Mutant alleles of MpRR-MYB5 but not MpRR-MYB2 appeared pale (**Figure 1D, E, F, Figure S1A-B**) and analysis of chlorophyll content confirmed this (**Figure 1H**). All lines in which insertions or deletions introduced premature stop codons in MpRR-MYB5 (**Figure S1A**) had 40-50% less chlorophyll than controls (**Figure 1E and H**). The Mp*rr-myb5* mutant was complemented when MpRR-MYB5 was expressed from its own promoter (**Figure S1C**) confirming that the pale phenotype was unlikely associated with off-target CRISPR/Cas9 editing. Mutating MpRR-MYB5 and MpRR-MYB2 simultaneously (**Figure S1D**) led to extremely pale plants with chlorophyll content reduced to 95% compared with controls (**Figure 1G and H**). To test whether the photosynthetic apparatus was functional in the single Mp*rr-myb5* and double Mp*rr-myb5,2* mutants we applied the inhibitor Di-Chlorophenyl Di-Methyl Urea (DCMU) that blocks photosynthetic electron transport^28^ and measured activity of photosystem II via chlorophyll fluorescence imaging. This showed that although these mutants had low levels of chlorophyll, the photosynthetic apparatus was operational ( **Figure 1I**). Consistent with the very low chlorophyll levels in Mp*rr-myb5*, and Mp*rr-myb5,2* double mutants, chloroplasts were significantly smaller and thylakoids underdeveloped (**Figure 1J-R and Figure S1E**). It was noticeable that poorly developed chloroplasts from double Mp*rr-myb5*,*2* mutants contained significant amounts of starch.

To determine whether MpRR-MYB5 and MpRR-MYB2 limit greening, we generated overexpression lines driven by the strong MpUBE2 constitutive promoter^29^ and to facilitate analysis of chloroplast size per cell used GFP to mark the plasma membrane (**Figure S2A-E**). Although quantitative polymerase chain reactions confirmed that each transgene was over-expressed (**Figure S2F-N**) plants appeared similar to controls and there were no evident perturbations to chlorophyll content, chloroplast size or morphology (**Figure S2O-P**). We conclude that MpRR-MYB5 and MpRR-MYB2 act redundantly and are necessary for chloroplast biogenesis but in contrast with Mp GLK^15^ they are not sufficient to activate this process. Moreover, in the absence of both Mp RR-MYB5 and MpRR-MYB2 assembly of the photosynthetic apparatus is very limited.

### MpRR-MYB5&2 act with MpGLK to control chloroplast biogenesis

As double Mp*rr-myb5*,*2* mutants showed residual chloroplast development and were viable, we hypothesised that the limited ability for photoautotrophic growth was associated with activity of the previously characterised master regulator GLK. To test this, we attempted to generate higher order mutants which combined mutant alleles of Mp*glk*, Mp*rr-myb5* and Mp*rr-myb2*. We were able to knock out MpRR-MYB5 in the presence of Mp*glk* mutant alleles (**Figure S3A-B**). Such double mutants were paler than the single Mp*glk* mutant (**Figure 2A-D**) and contained less chlorophyll (**Figure 2E**). Application of DCMU confirmed that the photosystem II was functional in the Mp*glk*,*rr-myb5* double mutant (**Figure 2F**). Double Mp*glk,rr-myb5* mutants had smaller chloroplasts with fewer thylakoid membranes and reduced granal stacking compared with each single mutant (**Figure 2G-O**). Thus, in the absence of both MpRR-MYB5 and MpGLK, very limited assembly of the photosynthetic apparatus takes place.

**Figure 2:**
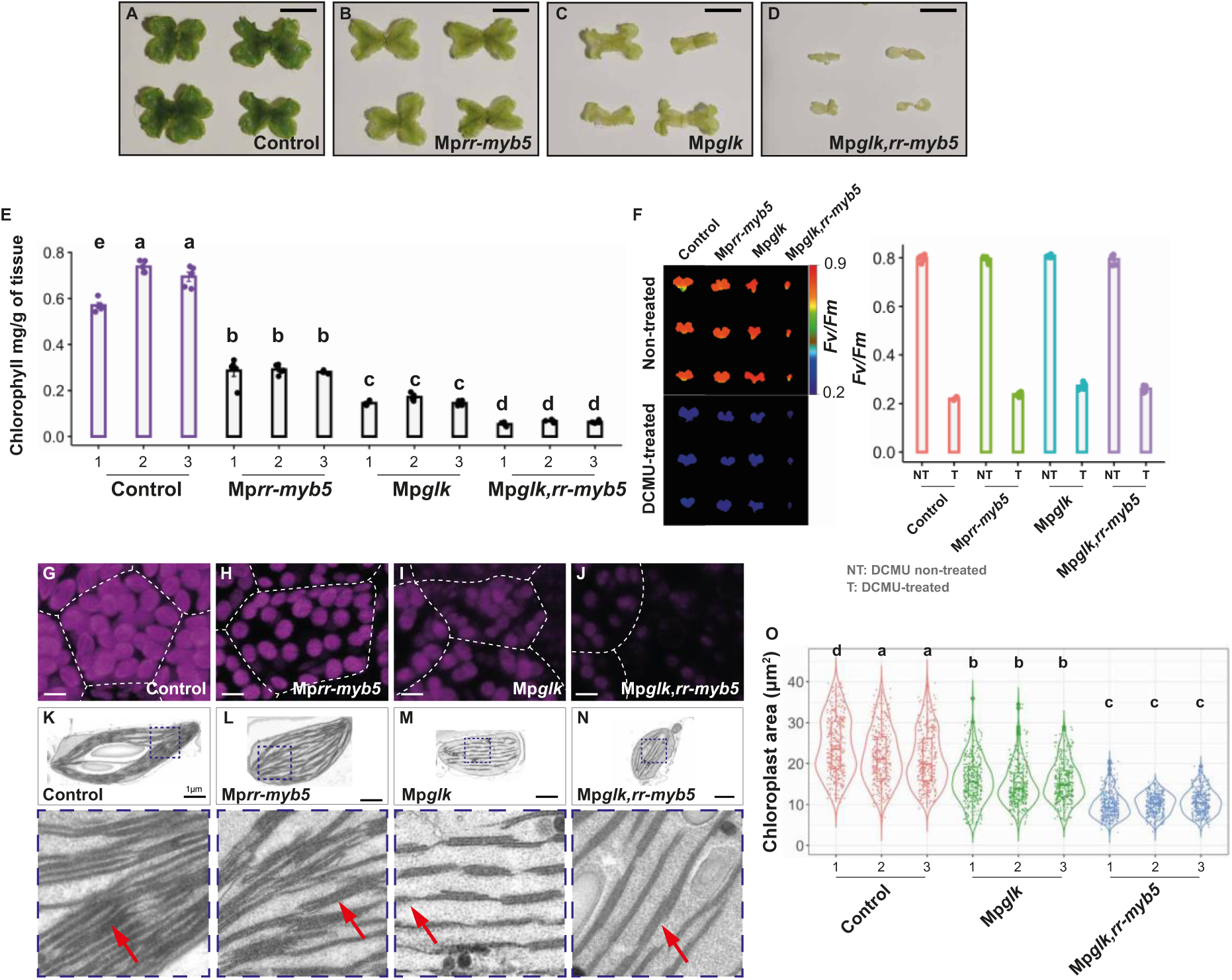
MpRR-MYB5 acts synergistically with MpGLK to control chloroplast biogenesis. **A-D)** Representative images of control, Mp*glk*, Mp*rr-myb5* and Mp*glk,rr-myb5* mutants. Scale bars represent 7 mm. **E)** Chlorophyll content is lower in Mp*glk*, Mp*rr-myb5* and the double Mp*glk,rr-myb5* mutants compared with controls. Letters show statistical ranking using a *post hoc* Tukey test with different letters indicating statistically significant differences at p<0.01. Values indicated by the same letter are not statistically different, n=5. **F)** Representative images and quantification after imaging of the chlorophyll fluorescence parameter *F_v_/F_m_* with and without the inhibitor Di-Chlorophenyl Di-methyl Urea (DCMU). Mp*rr-myb5,* Mp*glk* and Mp*glk,rr-myb5* mutants have lower chlorophyll content, the drop in *F_v_/F_m_* values after treatment with DCMU indicates that photosystem II is functional. **G-J)** Representative images after confocal laser scanning microscopy of control, Mp*rr-myb5,* Mp*glk* and Mp*glk,rr-myb5* mutants. Chlorophyll autofluorescence shown in magenta, cell borders marked with dashed white lines. Scale bars represent 10 μm. **K-N)** Representative images from transmission electron microscopy images of control, Mp*rr-myb5,* Mp*glk* and Mp*glk,rr-myb5* mutants. Scale bars represent 1 μm. The dashed area depicted in each chloroplast is enlarged and granal stacks indicated with red arrows. **O)** Chloroplast area of Mp*glk* and Mp*glk,rr-myb5* mutants. Box and whiskers represent the 25 to 75 percentile and minimum-maximum distributions of the data. Letters show statistical ranking using a *post hoc* Tukey test with different letters indicating statistically significant differences at p<0.01. Values indicated by the same letter are not statistically different. n=330.

We were unable to generate triple Mp*glk,rr-myb5,2* mutants implying that this allelic combination is lethal. For example, after super transforming Mp*glk,rr-myb5* double mutants with a vector allowing expression of the same guide RNA used to generate the single Mp*rr-myb2* mutants reported above, 91 lines were obtained. However, none were paler than the double Mp*glk,rr-myb5* mutant, and when genotyped 86 lines had no edits in MpRR-MYB2. Of the five lines that were edited in MpRR-MYB2 (as well as MpGLK and MpRR-MYB5) genotyping showed that the mutations had limited impact on the MpRR-MYB2 protein. For example, these edits altered one, two, three or seven amino acids in a poorly conserved region of the protein, and in all cases reading frame was maintained (**Figure S3C and D**). In contrast, when the original single Mp*rr-myb2* mutants were identified (**Figure 2F**) 50% of plants contained mutations that introduced early stop codons or disturbed the reading frame. We conclude that absence of all three proteins (MpGLK, MpRR-MYB5 and MpRR-MYB2) is lethal likely because chloroplast biogenesis is abolished.

### MpRR-MYB transcription factors regulate genes allowing carbon fixation, photorespiration and photosystem function

To provide insight into the types of genes regulated by MpRR-MYB5 and MpRR-MYB2 we performed RNA sequencing of overexpressing lines as well as single and multiple mutants. Overexpression of MpRR-MYB2 and MpRR-MYB5 led to the upregulation of 71 and 11 genes respectively (*padj*-value ≤0.01, LFC ≥ 1-fold) (**Figure S4A and C**) and there was limited overlap between these two datasets (**Figure 3A and Figure S4G and H**). This contrasts with overexpression of MpGLK that led to the upregulation of 492 genes (**Figure 3A** and ^15^).

**Figure 3:**
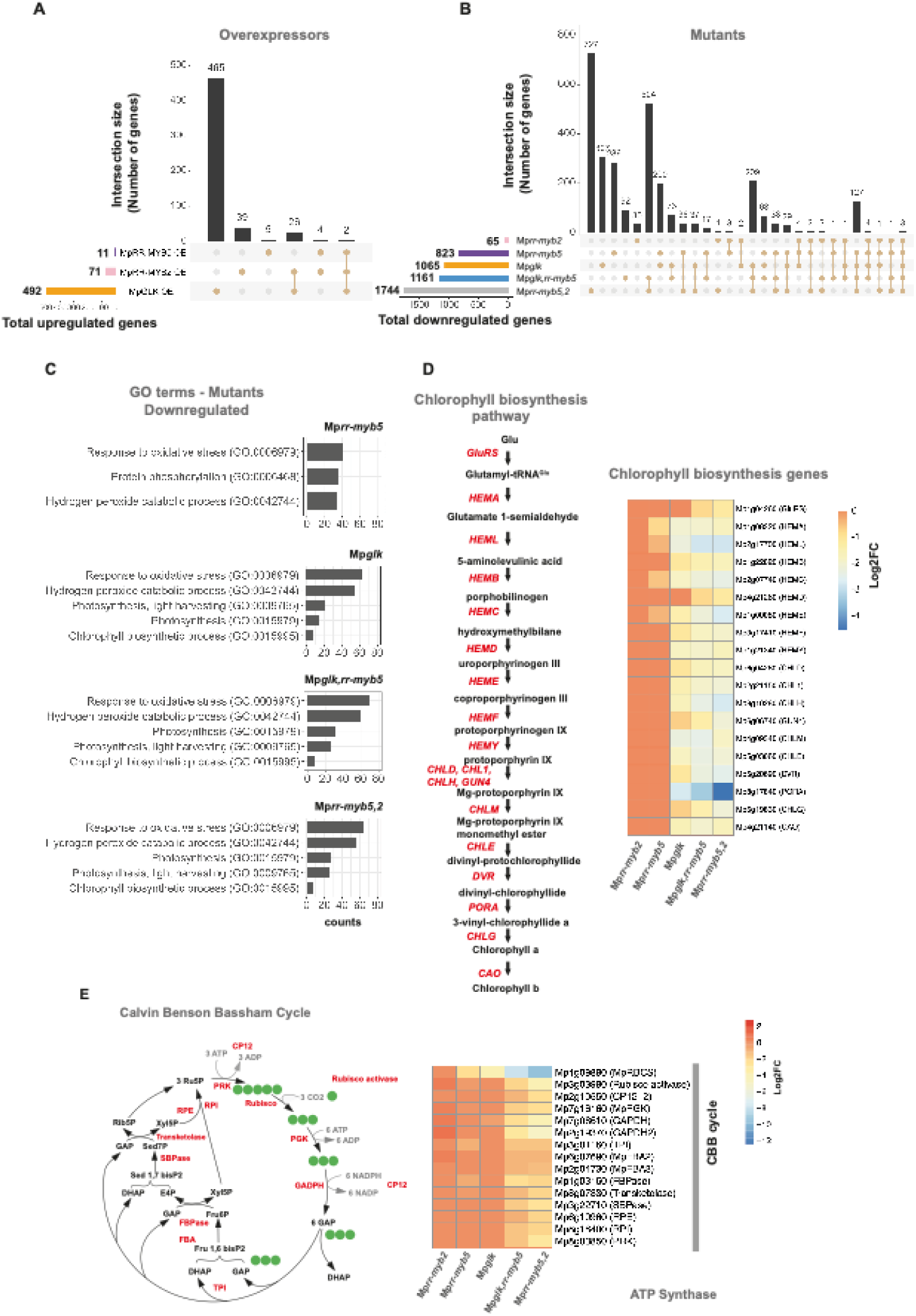
MpRR-MYB5 and MpRR-MYB2 regulate expression of genes encoding chlorophyll biosynthesis and the Calvin Benson Bassham cycle. **A)** Upset diagram showing sets of upregulated genes in MpRR-MYB5, MpRR-MYB2 and MpGLK over-expression lines. **B)** Upset diagram showing sets of downregulated genes in Mp*glk*, Mp*rr-myb2,* Mp*rr-myb5,* and the double Mp*glk,rr-myb5* or Mp*rr-myb5,2* mutants. **C)** Enriched Gene Ontology terms for Mp*rr-myb5,* Mp*glk,rr-myb5,* Mp*glk* and Mp*rr-myb5,2* mutants. **D)** Heatmap illustrating the extent of down-regulation of transcripts encoding enzymes of chlorophyll biosynthesis in Mp*rr-myb2,* Mp*rr-myb5, Mpglk,* and the *Mpglk,rr-myb5* or Mp*rr-myb5,2* mutant alleles. **E)** Heatmap indicating lower transcript abundance of genes encoding components of Calvin-Benson-Bassham cycle in Mp*glk*, Mp*rr-myb5,* Mp*rr-myb2* as well as Mp*glk,rr-myb5*,*myb2* and Mp*rr-myb5*,*myb2* double mutants. Modified from Lea and Leegood, 1998.

In loss of function mutants for MpRR-MYB2 or MpRR-MYB5, 65 and 823 genes, respectively, showed reductions in transcript abundance compared with controls (*padj*-value ≤0.01, LFC ≥ 1-fold) (**Figure 3B and Figure S4B and D**). Knocking out MpGLK had greater impact with 1065 genes being downregulated (**Figure 3B** and ^15^). In double Mp*glk,rr-myb5* mutants, 1161 genes had lower transcript abundance than controls, and in the double Mp*rr-myb5,2* mutants this was increased to 1744 (**Figure 3B**). The largest overlap in changes to transcript abundance between genotypes (524 genes) was detected for Mp*glk,rr-myb5* and Mp*rr-myb5,2* mutants (**Figure 3B**). This finding further supports synergistic action of Mp*RR-MYBs* and MpGLK. Gene Ontology (GO) terms were used to provide insight into the classes of genes impacted by overexpression or loss of function of Mp *RR-MYBs* and MpGLK. Consistent with the lack of detectable phenotype after overexpression of MpRR-MYB5 or MpRRMYB2, or loss of MpRR-MYB2 function, no distinct GO terms were impacted in these lines. Although the response to oxidative stress GO term was over-represented in both Mp*rr-myb5* and Mp*glk* mutants, other terms were distinct (**Figure 3C**). For example, Mp*rr-myb5* mutants showed changes to protein phosphorylation and peroxidase activity terms, whilst in Mp*glk* photosynthesis, light harvesting and chlorophyll biosynthesis terms were affected (**Figure 3C** and ^15^). It was notable that very similar GO terms responded in Mp*rr-myb5,2*, Mp*glk* and Mp*glk,rr-myb5* mutants (**Figure 3C**). For example, in all genotypes the top five biological processes impacted were response to oxidative stress, hydrogen peroxide catabolism, photosynthesis, light harvesting and chlorophyll biosynthesis (**Figure 3C**). Thus, loss of function alleles for Mp*rr-myb5,2,* Mp*glk* and Mp*glk,rr-myb5* all caused changes in GO terms primarily associated with photosynthesis.

Since chlorophyll content was reduced in Mp*rr-myb5* and Mp*rr-myb5,2* double mutants we examined impact on transcript abundance derived from genes associated with the nineteen annotated chlorophyll biosynthesis genes (**Figure 3D**). With the exception of the HEMA gene in Mp*rr-myb5* mutants, knocking out either MpRR-MYB5 or MpRR-MYB2 did not significantly affect transcript abundance from chlorophyll biosynthesis genes. In contrast, in Mp*glk* mutant alleles transcript abundance from seventeen chlorophyll biosynthesis genes was reduced, and in the Mp*rr-myb5,2* double mutant all nineteen genes were significantly downregulated (**Figure 3D**). We next examined the impact of loss of the Mp*RR-MYBs* on approximately 200 other genes annotated as photosynthesis related in *M. polymorpha* (**Dataset S1**). This group included genes associated with CO_2_ fixation and the light harvesting apparatus as well as their assembly and repair. In the single Mp*rr-myb5* and Mp*rr-myb2* mutant alleles there was limited effect on photosynthesis associated genes (**Figure 3E and Figure 4A**). For example, in Mp*rr-myb2* expression of only a single photosynthesis gene (petE Mp4g02720) was perturbed (**Dataset S2**). In Mp*rr-myb5* mutants, a small number of genes were impacted including those encoding a small subunit of RuBisCO (Mp4g09890) and a CHLOROPHYLL A/B BINDING PROTEIN (Mp7g05530) (**Figure 3E and Dataset S2**). As expected, changes to photosynthesis transcripts were more evident in the Mp*glk* mutant (**Figure 3E and Figure 4A**), and even more severe when both MpRR-MYB5 and MpGLK were mutated (**Figure 3E and Figure 4A**). Strikingly, when MpRR-MYB2 and MpRR-MYB5 were simultaneously knocked out, the effect on photosynthesis associated genes was extensive and more widespread than in the *Mpglk,rr-myb5* double mutant. For example, in Mp*rr-myb5,2* double mutants, the majority of genes encoding enzymes involved in the Calvin Benson Bassham cycle and photorespiration were downregulated (**Figure 3E and Figure S3I**). Moreover, genes encoding components of both photosystems and their respective light harvesting complexes as well as the Cytochrome *b_6_f* complex were downregulated (**Figure 4A**). We also found that genes associated with assembly of RuBisCO, non-photochemical quenching, as well as granal stacking and repair of photosystem II were impacted in Mp*rr-myb5,2* double mutants (**Figure S3J**). This contrasts with Mp*glk* in which only genes encoding enzymes of chlorophyll biosynthesis as well as components of the photosystems and their light harvesting complexes were mis-regulated (**Figure 4A**).

**Figure 4:**
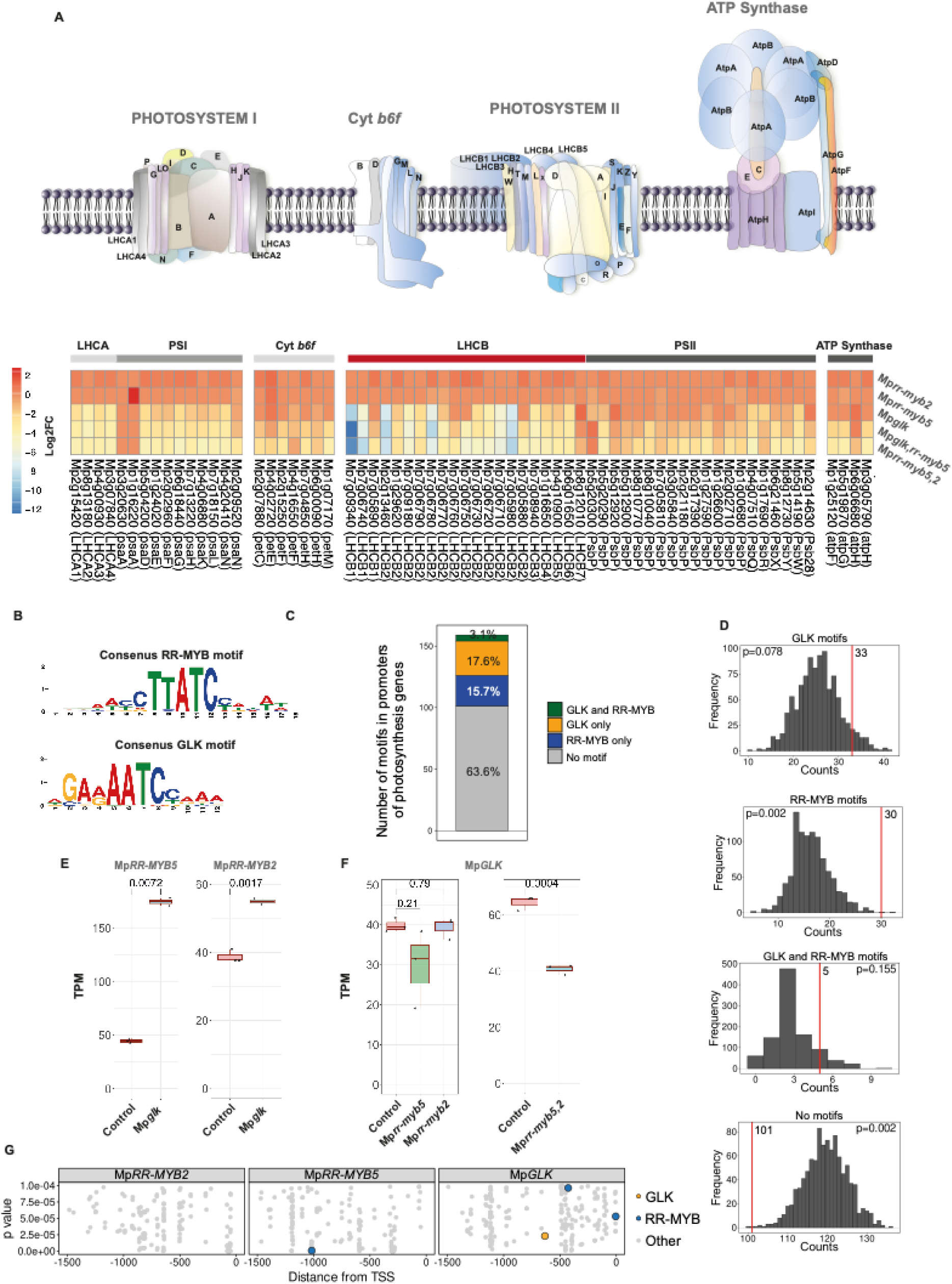
Combined loss of MpRR-MYB5&2 has greater impact on abundance of transcripts encoding the light harvesting apparatus than loss of MpGLK and down-regulates GLK expression. **A)** Heatmap indicating lower transcript abundance of genes encoding components of Photosystem II, Photosystem I and the cytochrome *b_6_f* complex in Mp*glk*, Mp*rr-myb5,* Mp*rr-myb2* as well as Mp*glk,rr-myb5*,*myb2* and Mp*rr-myb5*,*myb2* double mutants. The greatest degree of down regulation was detected in the double Mp*rr-myb5,2* double mutant. Schematic modified from^10^ and^25^. **B)** Consensus motif sequence logo for RR-MYB and GLK transcription factors. **C)** Bar chart showing RR-MYB and GLK binding sites within 500bp upstream of the predicted transcriptional start site (TSS) in *M. polymorpha* photosynthesis genes. **D)** Histograms showing the distribution of motifs in 500bp promoters of 1000 random gene sets. Red line indicates the frequency of the motif in *M. polymorpha* photosynthesis genes. P-values calculated by permutation testing. **E-F)** Transcripts abundance of MpGLK, MpRR-MYB2 and MpRR-MYB5 in control, Mp*rr-glk,* Mp*rr-myb5,* Mp*rr-myb2* and Mp*rr-myb5*,*2* mutant backgrounds. Data presented as Transcript Per Million (TPM), and P-values of two-tailed *t*-test are shown. **G)** Scatter plots showing position and predicted binding affinity of GLK, RR-MYB binding and other motifs upstream of MpGLK, MpRR-MYB2 and MpRR-MYB5 TSS, the y-axis shows p-values of matches between upstream regions and motif position weight matrices, and x-axis shows position of the motif center relative to the TSS. P-values calculated by log-likelihood score using FIMO.

### MpRR-MYB transcription factors condition expression of GLK

Consistent with the alterations in transcript abundance described above, consensus binding sites for the GLK and RR-MYBs proteins derived from ChIP-seq data^25^ and DAP-seq^30^ were found in promoters of 37% of photosynthesis genes (**Figure 4B**). 18% and 16% possessed motifs associated with GLK and RR-MYB binding respectively, and 3% contained both motifs (**Figure 4C and Dataset S3**). To test whether GLK and RR-MYB binding sites were enriched in photosynthesis genes above that expected by chance, we determined the frequency of these motifs in 500 base pairs upstream of 159 *M. polymorpha* photosynthesis genes compared with 1000 random sets of 159 *M. polymorpha* promoters (**Figure 4D**). 500bp was selected in order to focus analysis on core promoters rather than long distance enhancer elements and to reduce background signal associated with the increased probability of finding any motif by chance as the search space is greater (**Table S1**). Far fewer photosynthesis genes contained neither motif than would be expected from the background (p-value=0.002) whilst RR-MYB motifs were strongly enriched in photosynthesis genes (p-value=0.002). GLK motifs were over-represented compared with most background sets (p-value=0.078) although this enrichment was weaker than that for RR-MYB motifs. Overall, these data are consistent with overlapping as well as distinct roles for the two classes of transcription factor. Our data also support the notion that in contrast with MpGLK, the Mp*RR-MYBs* activate genes allowing CO_2_ fixation as well as light harvesting.

To better understand this interplay between MpRR-MYB2, MpRR-MYB5 and MpGLK we first examined the expression of MpRR-MYB2&5 in the *Mpglk* mutant background. MpRR-MYB5 and MpRR-MYB2 were upregulated in Mp*glk* (**Figure 4E**), a finding consistent with MpGLK acting to repress MpRR-MYB5&2. In contrast, in the Mp*rr-myb5,2* double mutant MpGLK was downregulated (**Figure 4F**). We were not able to identify any strong binding motifs for GLK in either MpRR-MYB5 or MpRR-MYB2 but in the promoter of MpGLK consensus binding sites for *RR-MYBs* were clearly evident (**Figure 4G**). The promoters of MpRR-MYB5 and MpGLK each contained their own binding site indicating the potential for self-regulation.

We next tested the extent to which MpRR-MYB2, MpRR-MYB5 or MpGLK transcription factors could rescue the pale phenotype of single Mp*rr-myb5,* Mp*glk,* and double Mp*glk,rr-myb5* or Mp*rr-myb5,2* mutants (**Figure 5A-F**). Quantitative polymerase chain reactions confirmed that each transgene was over-expressed (**Figure S5**). Both MpRR-MYB5 and MpRR-MYB2 complemented Mp*rr-myb5* mutants (**Figure 5B, C and G**), further arguing for functional redundancy between MpRR-MYB5 and MpRR-MYB2. However, neither MpRR-MYB rescued single Mp*glk* or double Mp*glk,rr-myb5* mutants (**Figure 5B, D, E and G**). When MpGLK was expressed in the Mp*rr-myb5,2* double mutant background chlorophyll levels were increased by ∼10% but the absolute levels were still 90% lower than wild type (**Figure 5F and H**). Consistent with functional redundancy between MpRR-MYB5 and MpRR-MYB2 we also found that MpGLK partially complemented Mp*rr-myb5* single mutants **Figure 5C and G**) since chlorophyll accumulation was lower compared to MpGLK overexpression in Mp*glk* mutants.

**Figure 5:**
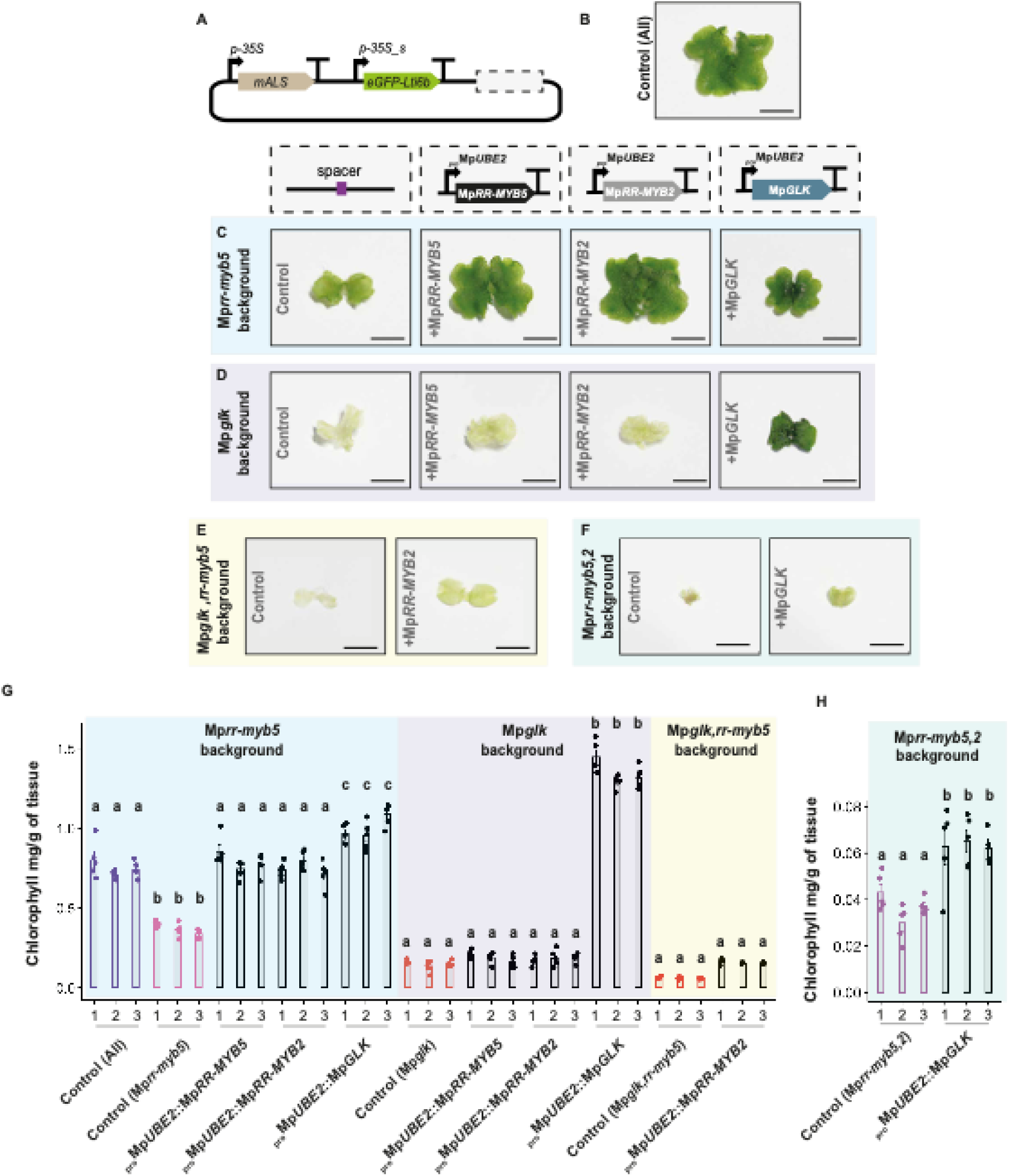
*M. polymorpha* RR-MYBs do not complement MpGLK: **A)** Schematic representation of constructs used to overexpress MpRR-MYB5, MpRR-MYB2 and MpGLK. **B)** Representative image of control plants. Scale bar represents 5 mm. **C-F)** Representative images of Mp*rr-myb5,* Mp*glk,* Mp*glk,rr-myb5* and Mp*rr-myb5,2* mutants complemented with MpRR-MYB5, MpRR-MYB2 and MpGLK. Scale bars represent 2mm. **G-H)** Chlorophyll content in control, Mp*rr-myb5,* Mp*glk,* Mp*glk,rr-myb5* and Mp*rr-myb5,2* mutants complemented with MpRR-MYB5, MpRR-MYB2 and MpGLK. Letters show statistical ranking using a *post hoc* Tukey test (comparisons are made between groups highlighted with the same colour rectangle, different letters indicating statistically significant differences at P<0.01). Values indicated by the same letter are not statistically different n=5.

### A conserved role for RR-MYBs in *Arabidopsis thaliana*

The RR-MYB/CCA1-like subfamily of MYB-related transcription factors^27,31,32^ containing MpRR-MYB5 and MpRR-MYB2 are characterised by a conserved SHAQK(Y/F)F motif (**Figure S6A**). Based on phylogenetic analysis we identified eleven members of this group in *A. thaliana* (**Figure 6A, Figure S6B-E**) of which AtMYBS1, AtMYBS2 and AT5G23650 were the closest homologs of MpRR-MYB5 and MpRR-MYB2 (**Figure S6B-E**). Re-analysis of publicly available data indicated that AT5G23650 is not expressed in photosynthetic tissues and so we focused analysis on AtMYBS1 and AtMYBS2. Due to the functional redundancy evident for MpRR-MYB5 and MpRR-MYB2 above, double At*mybs1,mybs2* mutants were identified after CRISPR/Cas9-mediated gene editing (**Figure S7A and B**) and analysed in parallel with previously generated single At*mybs1* and At*mybs2* mutants. There were no detectable changes to rosette phenotype in the single mutants but double At*mybs1,mybs2* mutants were pale (**Figure 6C-F**), and this was most noticeable after bolting (**Figure 6G**). Confocal laser scanning microscopy revealed no detectable changes in chloroplast size or number in mesophyll cells of single mutants, but chloroplasts were smaller in the double At*mybs1,2* mutant (**Figure 6F**). Notably, chloroplasts of At*mybs1,2* mutants contained underdeveloped thylakoids (**Figure 6H**) similar to Mp*rr-myb5,2* mutants. In support of these findings, chlorophyll content in the single mutants was indistinguishable from wild-type, but was ∼40% lower in the double At*mybs1,mybs2* mutant (**Figure 6Ι)**, and quantification of chloroplast size demonstrated a 50% reduction in the double mutant (**Figure 6J**).

**Figure 6:**
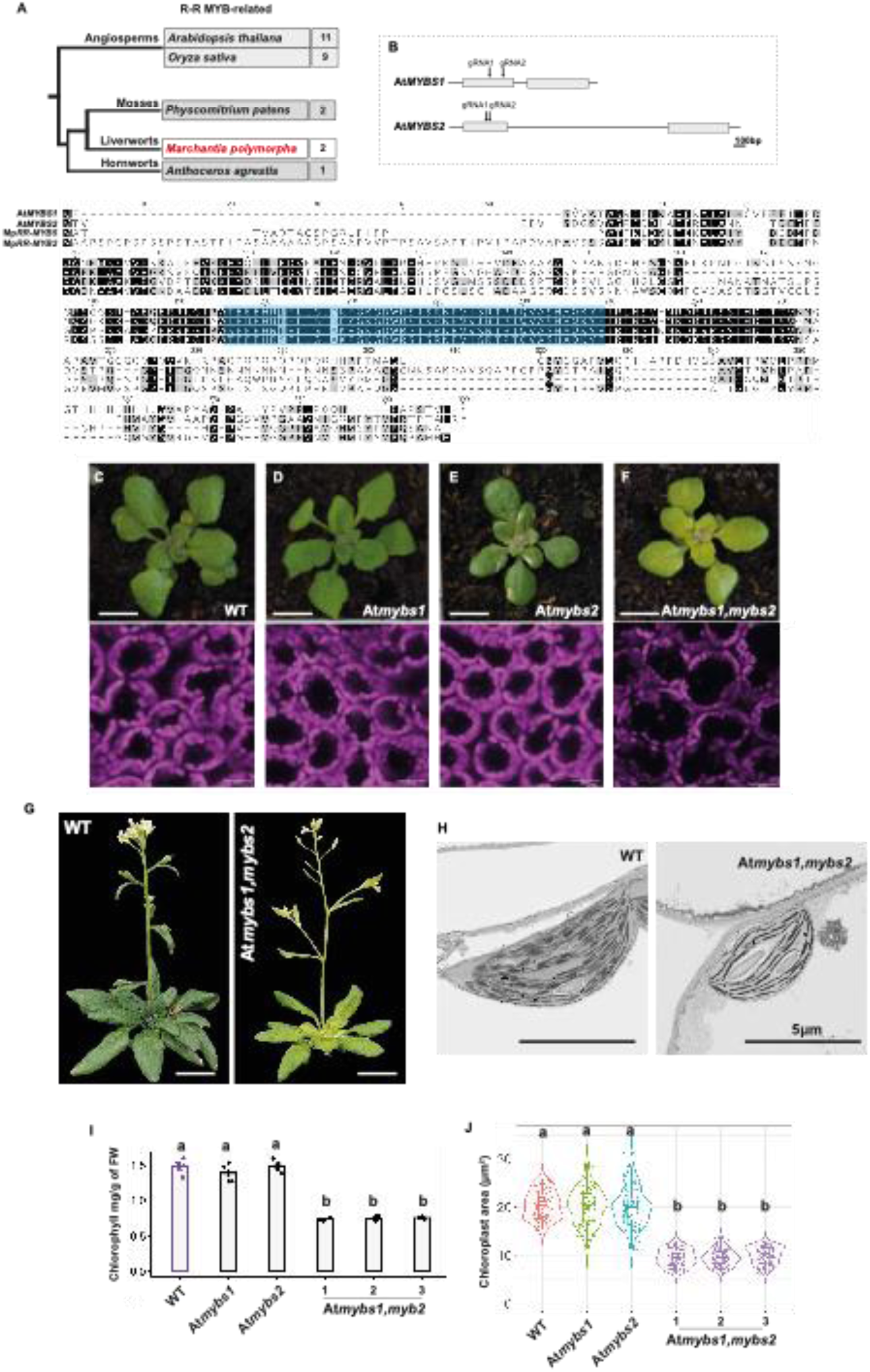
RR-MYBs control chlorophyll biogenesis in *A. thaliana.* **A)** Number of RR-Myb-Related transcription factors in *A. thaliana*, rice and bryophytes. Amino acid alignment of the two MpRR-MYB factors in *M. polymorpha* (red arrowheads) and *A. thaliana* MYBS1 and MYBS2 (bottom). Characteristic gene domains are shown as coloured boxes. **B)** Schematic representation of AtMYBS1 and AtMYBS2 gene structure showing exons as grey rectangles. guide (g) RNAs positions for gene editing shown with arrows. **C-F)** Images of two-week old seedlings of wild type, At*mybs1,* At*mybs2* and At*mybs1,mybs2* mutants (scale bars represent 7 mm. Representative images after confocal laser scanning microscopy also shown (bottom) with scale bars representing 25 μm. **G)** Images of wild type and At*mybs1,mybs2* mutants with inflorescence. Scale bars represent 1.5 cm. **H)** Representative transmission electron microscopy images of wild type and At*mybs1,mybs2* mutants. Scale bars represent 5 μm. **Ι)** Bar plots of chlorophyll content for wild type, At*mybs1,* At*mybs2* and At*mybs1,mybs2* mutants (three independent lines were used for measurements). Letters show statistical ranking using a *post hoc* Tukey test (with different letters indicating statistically significant differences at P<0.01). Values indicated by the same letter are not statistically different, n=5. **J)** Violin plots of chloroplast area for wild type, At*mybs1,* At*mybs2* and At*mybs1,mybs2* mutants. Box and whiskers represent the 25 to 75 percentile and minimum-maximum distributions of the data. Letters show statistical ranking using a *post hoc* Tukey test (with different letters indicating statistically significant differences at P<0.01). Values indicated by the same letter are not statistically different. n=100.

To gain insight into the types of photosynthesis genes regulated by AtMYBS1 and AtMYBS2 we performed RNA sequencing of the double mutant. 841 genes were down-regulated (*padj*-value ≤0.01, LFC ≥ 1-fold) and GO term analysis indicated that the two top biological terms impacted were response to light stimulus and photosynthesis (**Figure S7C and D**). Thus, consistent with lower levels of chlorophyll, loss of MYBS1 and MYBS2 in *A. thaliana* led to down-regulation of transcripts associated with photosynthesis. DNA motifs bound by GLK and the RR-MYBs have been determined.^25,30^ Promoters of the AtMYBS1&2 transcription factors in *A. thaliana* contained motifs recognised by GLK as well as RR-MYBs (**Figure 6A**) implying regulatory interplay. Although the promoter of GLK1 in *A. thaliana* has a motif associated with GLK binding (**Figure 7A**), in contrast with *M. polymorpha,* neither AtGLK1 or AtGLK2 contained any binding sites for MYBS1 or MYBS2 (**Figure 7A**). Promoters of the CGA1 and GNC transcription factors that regulate photosynthesis gene expression in *A. thaliana*^9,33^ also contained GLK but not MYBS predicted binding sites (**Figure 7A**). Analysis of GLK, CGA1 and GNC transcript abundance in the At*mybs1,2* indicated no change in GLK1 and CGA1 but increases in GLK2 and GNC (**Figure 7B**). This implies that cryptic or more distant MYBS1&2 binding sites exist for GLK2 and GNC, or that indirect regulation links expression of these transcription factors.

**Figure 7:**
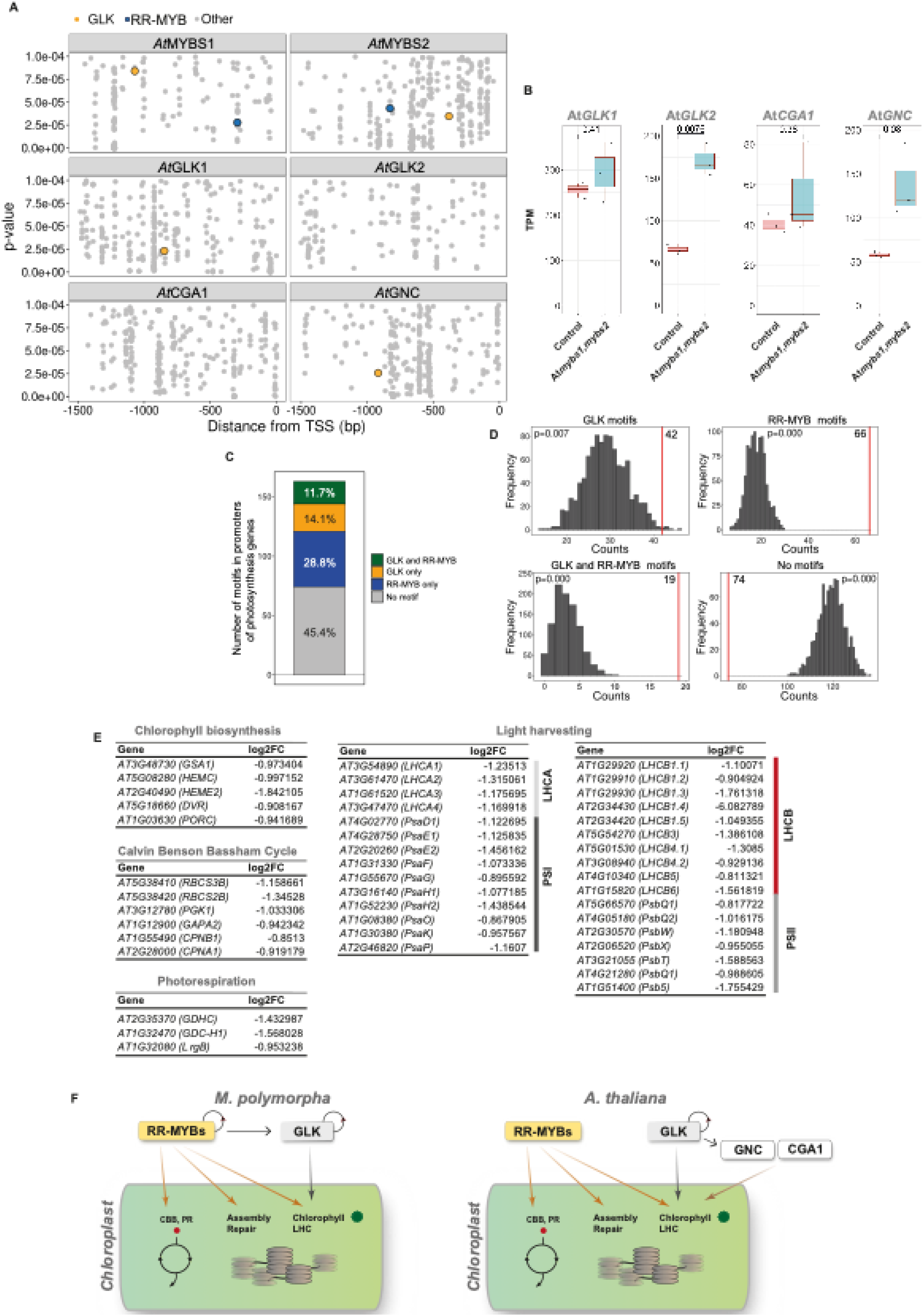
Loss of AtMYBS1&2 downregulates expression of genes encoding chlorophyll biosynthesis enzymes, the Calvin Benson Bassham cycle as well as those encoding the light harvesting apparatus. **A)** Scatter plots showing position and predicted binding affinity of GLK, RR-MYB binding and other motifs upstream of AtGLK1, AtGLK2, AtMYBS1 and AtMYBS2 transcriptional start site (TSS). Y-axis shows p-values of matches between upstream regions and motif position weight matrices, and x-axis shows position of the motif center relative to the TSS. P-values calculated by log-likelihood score using FIMO. **B)** Transcript abundance in Transcripts Per Million (TPM) of AtGLK1, AtGLK2, AtCGA1 and AtGNC in wild-type and At*mybs1,mybs2* mutant background. P-values of two-tailed *t*-test are shown. **C)** Barchart showing GLK and RR-MYB binding sites within 500bp upstream of the transcriptional start site (TSS) in *A. thaliana* photosynthesis genes **D)** Histograms showing the distribution of motifs in 500bp promoters of 1000 random gene sets. Red line indicates the frequency of the motif in *A.thaliana* photosynthesis genes. P-values calculated by permutation testing. **E)** List of Chlorophyll biosynthesis, Calvin Benson Bassham (CBB) cycle, Photorespiration and Light harvesting genes downregulated in At*mybs1,mybs2* mutants. **F)** Model illustrating role of RR-MYBs in the control of chloroplast biogenesis. In the bryophyte *M. polymorpha* (left), RR-MYBs act upstream of GLK. Both MpRR-MYB and MpGLK appear to regulate themselves through feedforward loops. Whilst MpGLK regulates photosynthesis genes associated with chlorophyll biosynthesis and sub-units of the light harvesting complexes (LHC), RR-MYBs also regulate genes of the Calvin Benson Bassham (CBB) cycle and photorespiration (PR) as well as assembly and repair of the photosystems. In the angiosperm model *A. thaliana* (right) RR-MYBs no longer act upstream of GLK but they do regulate genes of the Calvin Benson Bassham (CBB) cycle and photorespiration (PR) as well as assembly and repair of the photosystems. AtGLK regulates itself but also AtGNC, and as with *M. polymorpha*, it controls photosynthesis genes associated with chlorophyll biosynthesis and sub-units of the light harvesting complexes (LHC).

Despite these changes to the upstream regulatory network compared with *M. polymoprha*, multiple lines of evidence point to a role for AtMYBS1&2 in the control of photosynthesis gene expression in *A. thaliana*. For example, 500 base pair promoters from 29% and 14% of photosynthesis associated genes contain MYBS1&2 and GLK binding sites respectively (**Figure 7C and Dataset S3**) and 12% contained both (**Figure 7C and Dataset S3**). As with *M. polymorpha*, permutation analysis indicated that substantially fewer photosynthesis genes contained neither motif (p-value=0.000) (**Figure 7D**). Rather, they contained significantly more GLK, RR-MYB and both GLK and RR-MYB binding sites (p-value=0.007, 0.000 and 0.000 respectively) than would be expected by chance (**Figure 7D**). Moreover, abundance of transcripts associated with chlorophyll biosynthesis, both photosystems and their light harvesting apparatus, as well as the Calvin Benson Bassham cycle were perturbed when AtMYBS1&2 were knocked out in *A. thaliana* (**Figure 7E and Figure S7E**).

## Discussion

Chloroplasts allow photosynthesis, nitrogen and sulphur assimilation, as well as the biosynthesis of amino acids, fatty acids and carotenoids and so understanding their biogenesis has long been of interest.^34–38^ Moreover, it is now widely recognised that photosynthesis has not been optimised by evolution, and so a targeted reengineering of the process could contribute to crop development.^6,7^ Indeed, improvements in yield have been reported after over-expression of SEDOHEPTULOSE BISPHOSPHATASE,^39,40^ faster relaxation of non-photochemical quenching of photosystem II^41^ and rerouting of photorespiration.^42^ A complementary approach to improving photosynthesis predicted to increase yield by up to 50% would be to convert C_3_ crops to use the more efficient C_4_ pathway.^43,44^ Introducing C_4_ photosynthesis in C_3_ crops such as rice would require a remodelling of chloroplast biogenesis in mesophyll and bundle sheath cells.^45^

In land plants the GLK family of transcription factors are master regulators of chloroplast biogenesis, and CGA1 and GNC from the GATA family are considered ancillary players.^8,10,33^ Overexpression of GLK in rice is sufficient to increase chloroplast occupancy of cells such as the bundle sheath, and thus to partially phenocopy traits associated with the efficient C_4_ pathway.^36^ However, we have an incomplete understanding of transcription factors allowing chloroplast development. For example, GLK and CGA1/GNC loss of function mutants still possess small chloroplasts^9^ indicating that either these mutants are hypomorphic, or that additional unidentified actors control chloroplast biogenesis. Here we report two RR-MYB related transcription factors that act redundantly to control chlorophyll biosynthesis and photosynthesis associated gene expression in the bryophyte *M. polymorpha*. Homologs control chloroplast biogenesis in *A. thaliana* indicating functional conservation between these distantly related species. Interestingly, we were unable to identify null mutants lacking both GLK and RR-MYB in *M. polymorpha*. Indeed, although we used super-transformation of existing mutant alleles when attempting to generate triple mutants we did not observe white sectors. This apparent lethality of triple Mp*glk,rr-myb5,2* mutants mirrors loss-of-function mutations in plastidial pathways such as amino acid, vitamin, nucleotide or fatty acid biosynthesis, and those involved in chloroplast protein translation that result in an arrest of embryo development in *A. thaliana.*^46,47^ This often appears to coincide with the globular-to-heart transition stage when chloroplasts start to differentiate. Mutants in genes encoding plastidial proteins required for import, modification and localisation of indispensable proteins in the chloroplast are also often associated with embryo lethality.^46,47^ It is therefore possible that plants lacking both GLK and the RR-MYBs are unable to differentiate chloroplasts from proplastids.

The precise architecture of the gene regulatory network involving the *RR-MYBs* and GLK will need to be fully elucidated. However, consistent with the importance of negative feedback loops in plant development^48,49^ RR-MYBs induce GLK and GLK then represses the *RR-MYBs*. Transcripts from both Mp*RR-MYBs* were more abundant in the Mp*glk* mutant, and *MpGLK* contained predicted binding sites for the MpRR-MYBs. In Mp*rr-myb2,5* mutant alleles, transcripts derived from MpGLK were less abundant. Moreover, when the MpRR-MYBs were overexpressed in the Mp*glk* mutant it remained pale, arguing against RR-MYBs acting downstream of GLK. The stronger and broader down regulation of photosynthesis transcripts in the double Mp*rr-myb5,2* mutant compared with Mp*glk* could be due to the RR-MYBs acting upstream of GLK and other players. Alternatively, the larger response of photosynthesis transcripts in Mp*rr-myb5,2* mutants compared with Mp*glk* could be because the RR-MYBs have a broader set of targets. Although MpRR-MYB5 overexpression failed to upregulate MpGLK transcripts, MYB transcription factors commonly act in multimeric complexes involving bHLH and WD40 proteins^50^ or with MYCs proteins.^51^ It is therefore possible that additional partners need to be over-expressed in combination with the RR-MYBs to increase expression of MpGLK. It is also well documented that GLK is subject to multiple levels of regulation that can be overcome when non-native versions of the gene are mis-expressed.^36^ Thus, it may be that the lack of response in MpGLK after over-expression of MpRR-MYBs is due to a similar regulatory system. When MpGLK was mis-expressed in the double Mp*rr-myb5,2* mutant although chlorophyll content remained low (90% of wild type) it was significantly statistically increased. We interpret these data in two ways. Either MpGLK is downstream of the Mp*RR-MYBs* and both classes of transcription factor are needed to drive full photosynthesis gene expression, or MpGLK is permissive for very early stages of chloroplast biogenesis, but full assembly of the photosynthetic apparatus is strengthened by RR-MYBs because they positively regulate MpGLK and they also control genes allowing carbon fixation. A permissive role for GLK in initiating chloroplast biogenesis is consistent with its ability to convert cell types accumulating a small to a large chloroplast compartment.^36^ Based on the above, our current favoured hypothesis is that the RR-MYBs condition expression of GLK to permit early stages of chloroplast biogenesis. Although this conditioning can be partially overcome by over-expression of MpGLK in the presence of RR-MYBs^15^ in the absence of Mp*RR-MYBs* the impact of MpGLK over-expression is limited.

As would be expected from the evolutionary distance, rewiring has taken place between these transcription factors and the structural photosynthesis genes they target in *A. thaliana* and *M. polymorpha*. In *M. polymorpha* both the RR-MYB and GLK appear to regulate themselves, and the RR-MYBs condition expression of GLK **(Figure 7E**). In *A. thaliana* this regulatory system is more complex. For example, regulation of AtGLK by AtMYBS1&2 is less evident, but members of GATAs^9,33^ also control photosynthesis gene expression. Moreover, inducible overexpression of AtMYBS1 in *A. thaliana* has been reported to increase expression of photosynthesis genes^24^. Although similar sets of genes were down-regulated in loss of function Mp*rr-myb5,2* mutants, we did not detect widespread upregulation of photosynthesis genes after over-expression of *M. polymorpha* RR-MYBs. Also consistent with rewiring between these species is the fact that compared with MpRR-MYB2, the pale phenotype of MpRR-MYB5 indicates it plays a dominant role in chloroplast biogenesis while in *A. thaliana* neither single mutant was pale. Lastly, changes to the structure of the gene regulatory network associated with photosynthesis is also supported by the fact that promoters of MpRR-MYB5&2 contain their own binding sites, but MpGLK does not. In contrast, in *A. thaliana* promoters of AtMYBS1 and AtMYBS2 possess *cis*-elements for both GLKs and the RR-MYBs. The output is that the RR-MYBs operate upstream of GLK in *M. polymorpha* but not in *A. thaliana*.

The role of RR-MYBs in controlling chloroplast biogenesis is in fact supported by previous work. For example, in tomato LeMYBI has been reported to bind the promoter of the RBCS gene^52^. And, although no effect on chloroplast biogenesis was reported, a reduction in RBCS and the *chlorophyll a/b binding protein 1* (CAB1) genes expression has been reported in At*mybs1* mutants^53^. Along with transcription factors belonging to the GLK, B-Box and Nuclear Factor-Y families, random forest analysis of gene expression recently predicted that RR-MYBs regulate photosynthesis gene expression.^24^ Moreover, RNA sequencing of an inducible AtMYBS1 over-expressor line showed upregulation of photosynthesis genes that we detected as downregulated in the At*mybs1,mybs2* mutant. Based on these findings we conclude that the RR-MYB class of transcription factors likely control photosynthesis in a wide range of land plants.

Penetrance of the RR-MYBs on photosynthesis gene expression and chloroplast biogenesis in *M. polymorpha* was striking, with Mp*RR-MYBs* activating genes allowing CO_2_ fixation as well as light harvesting. Chloroplasts of Mp*myb5,2* mutants were ∼30% smaller than those of Mp*glk* mutants, and double Mp*myb5,2* mutants were paler than those of Mp*glk*. This appears to be because RR-MYBs control an overlapping but broader set of photosynthesis genes than those downstream of GLK. Previous work supports the notion that these two classes of transcription factors have shared targets, as co-binding of RR-MYB and GLK to photosynthesis genes has been proposed.^54^ Such cooperative binding of transcription factors is thought to allow greater variety of expression outputs.^55,56^ The reach of the RR-MYBs appears extensive in that they control genes encoding enzymes of the Calvin Benson Bassham cycle and photorespiration but also assembly and repair of RuBisCO. It seems likely that the large number of genes encoding a wide range of components underpinning photosynthesis targeted by the RR-MYBs contributes to the severe perturbation to phenotype when their function is removed. And, when combined with loss of GLK, this may be why lethality ensues. Overall, the data are consistent with overlapping as well as distinct roles for these two classes of transcription factor.

In summary, from analysis of *M. polymorpha* and *A. thaliana* whose last common ancestor diverged around 400 million years ago we propose a model in which both RR-MYBs and GLKs operate as master regulators of photosynthesis gene expression (**Figure 7F**). In both species the RR-MYBs play a conserved role in controlling photosynthesis gene expression and their targets are broader than those documented for GLK. As RR-MYBs appear ubiquitous in land plants^27^ it seems plausible they play a conserved role in chloroplast biogenesis. While we were unable to detect MpRR-MYBs in the Zygnematophyceae algae that are sister to the land plants^57^ they are in fact present in the Klebsormidiophyceae and Charophyceae^58,59^ (**Figure S6B**) that represent the other two most closely related algal lineages to land plants. GLK homologs are present in green algae^57^ and both GLK and RR-MYB motifs are present in promoter regions of these genes in *K. flaccidum* and *C. braunii* (**Figure S7G**). These data imply that *RR-MYBs* operated alongside GLK to control chloroplast biogenesis before the colonisation of land by plants.

## Materials & Methods

### Plant growth and transformation

*Marchantia polymorpha* Cam-1 (male) and Cam-2 (female) were grown on half-strength Gamborg B5 medium plus vitamins (Duchefa Biochemie G0210, pH 5.8) and 1.2% (w/v) agar (Melford capsules, A20021) under continuous light at 22 °C with light intensity of 100 μmol m^−2^s^−1^. *Arabidopsis* thaliana C*ol-0* was except the At*mybs2* mutant that was in the *Col-3* background, and plants were grown on F2 soil (Levington, F20117800) under 16 hours light, 8 hours dark at 20 °C, 60% humidity, and 150 μmol m^−2^ s^−1^ light. *Arabidopsis* T-DNA insertion mutants At*mybs1* (SAIL_1184_D04) and At*mybs2* (SALK_150774) were obtained from NASC with T-DNA and zygosity confirmed by PCR (**Figure S7F and Table S2**). For gene editing guide RNAs were predicted using CasFinder tool (https://marchantia.info/tools/casfinder/). Several gRNAs (**Table S3**) per target were tested.^60^ Single gRNAs were cloned^29^ into the destination vector pMpGE013^61^. To complement Mp*rr-myb5* mutants, the MpRR-MYB5 promoter from^62^ was used. For overexpression MpRR-MYB2 and MpRR-MYB5 coding sequences were synthesised (Integrated DNA Technologies) and cloned into the pUAP4 vector.^29^ For complementation, guide RNA resistant MpGLK,MpRR-MYB2 and MpRR-MYB5 coding sequences were synthesised (Integrated DNA Technologies) and cloned into the pUAP4 vector. OpenPlant parts used are listed in Star Methods.

To generate *A. thaliana* MYBS1/MYBS2 mutants two gRNAs per gene were cloned into the pEn-Chimera vector^63^ using a modification of the gRNA-tRNA approach.^64^ This placed guides into the pRU294 vector that has a codon optimised and intron-containing version of *Cas9* (*zCas9i*) driven by the egg-cell specific *pEC1.2* promoter.^65^ *A. thaliana* was transformed by floral dipping^66^ and genotyping performed as reported previously.^67^ T2 plants with confirmed edits were analysed (**Figure S7A and B**). For thallus transformation 5 mL LB media were inoculated with 3-4 *Agrobacterium* colonies (GV3101: 50 μg/mL rifampicin, 25 μg/mL gentamicin) and the plasmid-specific selection antibiotic. The preculture was incubated at 28°C for 2 days at 110 rpm. 5 mL of 2 d *Agrobacterium* culture were centrifuged for 7 min at 2000 x *g*. The supernatant was removed and pellet re-suspended in 5 mL liquid KNOP (0.25g/L KH_2_PO_4,_ 25g/L KCl, 25g/L MgSO_4_ 7H_2_O, 100g/L Ca(NO_3_)_2_ 4H_2_O, 12.5mg FeSO_4_7H_2_O, 30mM MES and pH5.5) plus 1% (w/v) sucrose and 100 μM acetosyringone. The culture was then incubated with shaking (120 rpm) at 28°C for 3-4 hours. Around 100 gemmae were transferred into a 6-well plate with 5 mL liquid KNOP medium supplemented with 1% (w/v) sucrose and 30 mM MES, pH 5.5, 80 μL of *Agrobacterium* culture and acetosyringone at final concentration of 100 μM. The tissue was co-cultivated with *Agrobacterium* for 3 days on a shaker at 110 rpm, at 22°C with ambient light. Using a sterile plastic pipette, the liquid was removed from each well and gemmae transferred onto plates with growth media containing the appropriate antibiotic (Chlorsulfuron 0.5 μM). To facilitate spreading of gemmae 1 -2 mL sterile water was added to the petri dish. To genotype *M. polymorpha* 3 × 3 mm pieces of thalli from individual plants were placed in 1.5 mL tubes and crushed in 100 μL genotyping buffer (100 mM Tris-HCl, 1 M KCl, 1 M KCl, and 10 mM EDTA, pH 9.5). Tubes were then placed at 70 °C for 15-20 mins and 380 μL sterile water added to each tube. 5 μL aliquots of the extract were used as a template for polymerase chain reactions.

### Chlorophyll determination, fluorescence measurements and imaging analysis

For chlorophyll measurements of *M. polymorpha* ∼30-50mg of 10-14 days old gemmalings were used with five biological replicates per genotype. Tissue was blotted dry before weighing and then transferred into a 1.5mL microfuge tube containing 1 mL of dimethyl sulfoxide (DMSO) (D8418, Sigma Aldrich) and incubated in the dark at 65 °C for 45 minutes. Samples were allowed to cool to room temperature for approximately one hour. Chlorophyll content was then measured using a NanoDrop™One/One C Microvolume UV-Vis Spectrophotometer (ThermoFisher) following the manufacturer’s instructions. Chlorophyll fluorescence measurements were carried out using a CF imager (Technologica Ltd, UK). *M. polymorpha* plants were placed in the dark for 20 mins and a minimum weak measuring light beam (<1 μmol m-^2^ s^-1^) applied to evaluate dark-adapted minimum fluorescence (F_o_), and a subsequent saturating pulse of 3000 μmol m^-2^ s^-1^ used to evaluate dark-adapted maximum fluorescence (*F_m_*). A total of three plants were measured per genotype and treatment. 20 µM DCMU (#45463, Sigma Aldrich) was added to half-strength MS media, and thalli placed in DCMU for 24 h before chlorophyll fluorescence measurements were obtained.

For confocal laser scanning microscopy, five to seven gemma were placed within a medium-filled gene frame together with 30 μL water prior to being sealed with a cover slip. Plants were imaged immediately using a Leica SP8X spectral fluorescent confocal microscope with either a 10X air objective (HC PL APO 10×/0.40 CS2) or 20X air objective (HC PL APO 20×/0.75 CS2). Excitation laser wavelength and captured emitted fluorescence wavelength window were 488 nm, 498−516 nm for GFP, and 488 or 515nm, 670−700 nm for chlorophyll autofluorescence. For electron microscopy ∼2 mm^2^ sections of 5-6 individual 3-week-old thalli were harvested, fixed, embedded and imaged as previously described.^68^ Chloroplast area was measured using ImageJ and the Macro in **Supplemental Information**.

### RNA extraction and sequencing

For *M. polymorpha*, RNA was extracted from 3-4 two-week old gemmae using the RNeasy Plant kit (#74903, Qiagen) with RLT buffer supplemented with beta-mercaptoethanol, and residual genomic DNA removed using the Turbo DNA-free kit (# AM1907, Invitrogen). 500 ng of DNase-treated RNA was used as template for cDNA preparation (SuperScript™ IV First-Strand Synthesis System, #18091050, Invitrogen) according to manufacturer’s instructions except that reverse transcription was 40 minutes and used oligo (dT)18 primers. qPCR was performed using the SYBR Green JumpStart *Taq* Ready Mix (#S4438, Sigma Aldrich) and a CFX384 RT System (Bio-Rad) thermal cycler. cDNA was diluted six times and oligonucleotides (**Table S3**) used at a final concentration of 0.5 μM. Reaction conditions comprised initial denaturation 94°C for 2 minutes followed by 40 cycles of 94°C for 15 seconds (denaturation) and 60°C for 1 minute (annealing, extension, and fluorescence reading). Primer sequences are in **Table S3**. Library preparation and RNA sequencing was performed by Novogene (Cambridge, UK). Briefly, messenger RNA was purified from total RNA using poly-T oligo-attached magnetic beads. After fragmentation, first strand cDNA was synthesised using random hexamer primers. Library concentration was measured on a Qubit instrument using the manufacturer’s procedure (Thermo Fisher Scientific) followed by real-time qPCR quantification. Library size distribution was analysed on a bioanalyzer (Agilent) following the manufacturer’s protocol. Quantified libraries were pooled and sequenced on a NovaSeq PE150 Illumina platform and 6G raw data per sample obtained. Adapter sequences were: 5’ Adapter: 5’- AGATCGGAAGAGCGTCGTGTAGGGAAAGAGTGTAGATCTCGGTGGTCGCCGTATCATT-3’. 3’ Adapter: 5’-GATCGGAAGAGCACACGTCTGAACTCCAGTCACGGATGACTATCTCGTATGCCGTCTTCTGCTTG-3’ FastQC was used to assess read quality and TrimGalore (https://doi.org/10.5281/zenodo.5127899) to remove low-quality reads and adapters. Reads were pseudo-aligned using Kallisto^69^ to the *M. polymorpha* genome version 5 (primary transcripts only, obtained from MarpolBase)^70^. Mapping statistics for each library are provided in **Supplemental Dataset 2**. DGE analysis was performed with DESeq2 ^71^, with padj-values < 0.01.

### Phylogenetic analysis

To identify RR Myb-related/CCA1-like genes three approaches were combined. Firstly, RR Myb-related/CCA1-like genes for twenty-one plant genomes were mined from iTAK^72^ and PlantTFDB^73^, Phytozome, Fernbase^74^, Phycozome and PhytoPlaza databases. Sequences for each species were aligned with the MpRR-MYB5 and *A. thaliana* MYBS1 and MYBS2 amino acid sequences using MAFFT^75^. Results were filtered manually to identify RR-Myb-related/CCA1-like orthologs distinguished from other Myb-related genes based on the conserved SHAKYF motif in the R1/2 domain. Secondly, BLASTP searches were performed against plant genomes in Phytozome v13, fern genomes (fernbase.org), hornworts genome (www.hornworts.uzh.ch)^76^, green algae genomes in PhycoCosm (/phycocosm.jgi.doe.gov), and 1KP using the MpRR-MYB5 and *A. thaliana* MYBS1 and MYBS2 amino acid amino acid sequence. Identified RR-Myb-related/CCA1-like protein sequences were aligned using MAFFT and trimmed using TrimAl^77^. A maximum likelihood phylogenetic tree was inferred using iQTree^78^, ModelFinder^79^ and ultrafast approximation for phylogenetic bootstrap^80^ and SH-aLRT test^81^. The tree was visualised using iTOL.^82^ Full list of sequences in **Dataset S4**.

### RR-MYB and GLK binding site analysis

AtGLK1 and GLK2 transcription factor binding motifs were taken from ChIP-seq data.^25^ AtMYBS1 and MYBS2 binding sites (motifs MA1186.1 and MA1399.1) were obtained from JASPAR.^83^ GLK and MYBS motifs were merged to create GLK combined and RR-MYB combined motifs and visualised using the Ceqlogo tool from MEME.^84^ The FIMO tool^85^ was used to scan promoter sequences of *A. thaliana* and *M. polymorpha* for matches to transcription factor binding motifs found in the JASPAR motif database.^83^ To account for input sequence composition, a background model was generated using the fasta-get-markov tool from the MEME suite.^84^ FIMO was then run with default parameters and a P value cut-off of 1 × 10^−4^. Matches to GLK and RR-MYB combined motifs were highlighted in each output. To assess occurrence of GLK and RR-MYB motifs in photosynthesis gene promoter sequences (500 bp upstream of the TSS) were scanned using FIMO^85^. Promoters were then scored for presence or absence of each motif and percentage of photosynthesis of genes containing each calculated. To test the background distribution of RR-MYB and GLK motifs a set of random promoters of the same size as the list of photosynthesis genes from *M. polymorpha* or *A. thaliana* was extracted and FIMO ran to determine the presence of these motifs. This process was iterated 1000 times and distributions of the frequency of these motifs plotted. The frequency of motifs in promoters of photosynthesis genes was also determined by searching each promoter with FIMO. Permutation tests were performed to test whether the frequency of motifs was significantly different to the frequency found in random selected promoters **(File S1).**

## Author contributions

N.Y.E., E.F. and J.M.H. designed the work. N.Y.E., E.F., K.B., T.S., M.T. and P.D. carried out the work. N.Y.E, E.F. and J.M.H. wrote the manuscript with input from all authors.

## Acknowledgements

This work was funded as part of the BBSRC/EPSRC OpenPlant Synthetic Biology Research Centre Grant BB/ L014130/1 to J.P.H., BBSRC BB/F011458/1 for confocal microscopy to J.P.H and BBSRC sLOLA BBP0031171 and ERACAPS grant C4BREED to J.M.H. T.B.S. was supported by a SNSF Postdoc Mobility Fellowship (P500PB_203128) and EMBO Long-Term Fellowship (ALTF 531-2019). For the purpose of open access, the authors have applied a Creative Commons Attribution (CC BY) licence to any Author Accepted Manuscript version arising from this submission. We thank Karin H. Müller, Filomena Gallo and Georgina E. Lindop from the Cambridge Advanced Imaging Centre for the electron microscopy sample preparation as well as the support during the image acquisition. We also thank Facundo Romani for useful comments and support during the project.

