## Supplemental Info for "Synergistic control of chloroplast biogenesis by *MYB-related* and *Golden2-like* transcription factors"

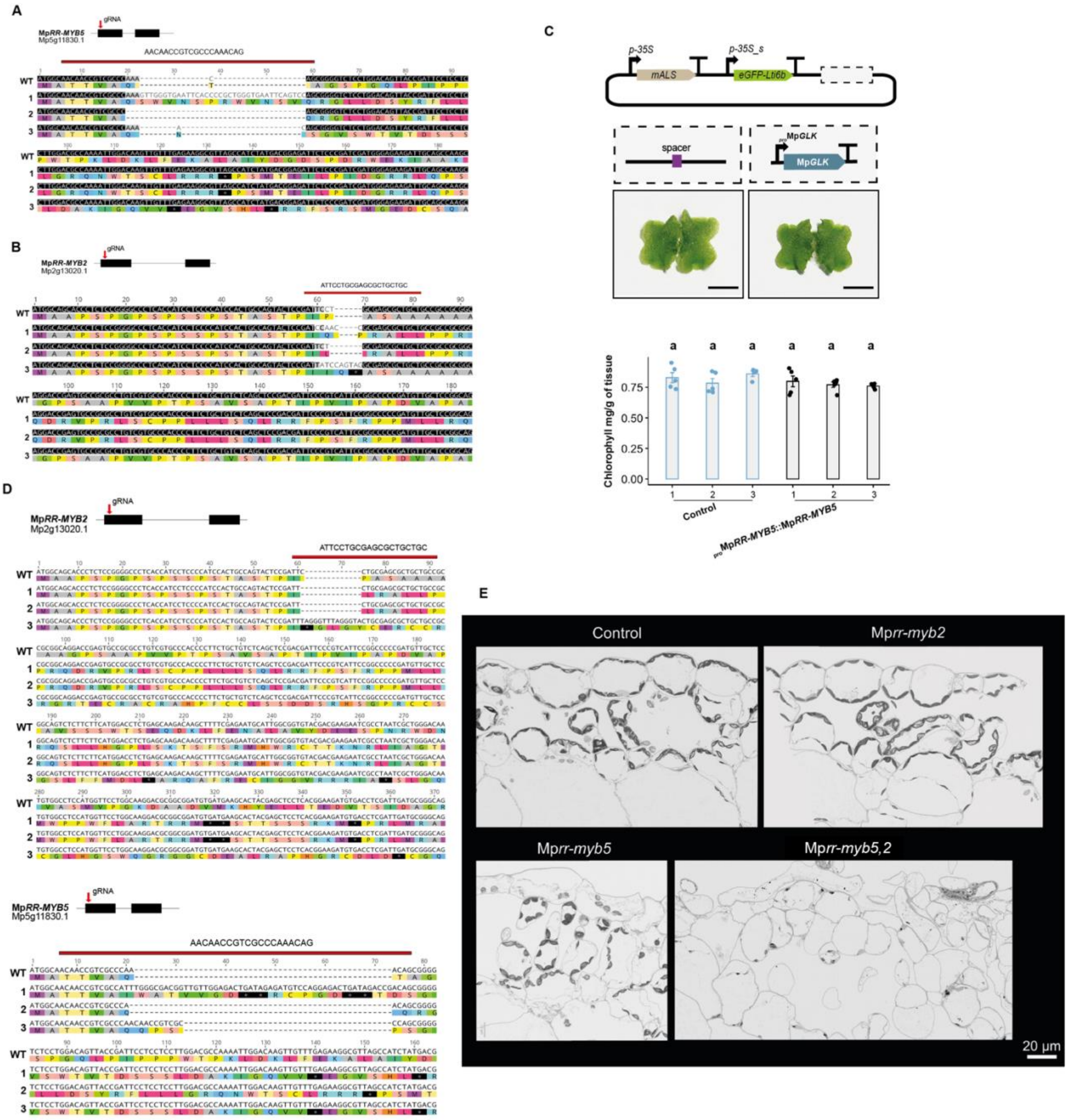
A
gRNA
MpRR-MYB5
Mp5g11830.1
AACAACCGTCGCCCAAACAG
B
MpRR-MYB2
Mp2g13020.1
ATTCCTGCGAGCGCTGCTGC
C
p-35S
p-35S_s
mALS
eGFP-Lti6b
spacer
proMpGLK
MpGLK
Chlorophyll mg/g of tissue
Control
proMpRR-MYB5::MpRR-MYB5
D
MpRR-MYB2
Mp2g13020.1
ATTCCTGCGAGCGCTGCTGC
MpRR-MYB5
Mp5g11830.1
AACAACCGTCGCCCAAACAG
E
Control
Mprr-myb2
Mprr-myb5
Mprr-myb5,2
20 µm

**Figure S1: CRISPR/Cas9 mediated gene editing and generation of deletions.**

**A)** Schematic representation of the Mp*RR-MYB5* gene structure showing exons as black boxes. Position of gRNA is shown with an arrow. Sequence analysis of Mp*rr-myb5* knockout mutant lines. The wild-type *Marchantia polymorpha* Cam-1 sequence is shown at the top, with the 20 bp gRNA target sequence highlighted with a red line. The amino acid sequence is depicted below the nucleotide sequence. **B)** Schematic representation of the Mp*RR-MYB2* gene structure showing exons as black boxes. Position of gRNA is shown with an arrow. Sequence analysis of Mp*rr-myb2* knockout mutant lines. The wild-type *M. polymorpha* Cam-1 sequence is shown at the top, with the 20 bp gRNA target sequence highlighted with a red line. The amino acid sequence is depicted below the nucleotide sequence. **C)** Top: Schematic representation of control construct and construct used to express Mp*RR-MYB5* using a 3-kilobase fragments upstream the ATG start codons as Mp*RR-MYB5* 'native' promoter regions. Bottom: phenotypes of control plants and Mp*rr-myb5* mutants complemented with the $_{pro}$Mp*RR-MYB5*::Mp*RR-MYB5.* Scale bars represent 5 mm. Chlorophyll content in control, Mp*rr-myb5* mutants complemented with the $_{pro}$Mp*RR-MYB5*::Mp*RR-MYB5.* Letters show statistical ranking using a *post hoc* Tukey test (with different letters indicating statistically significant differences at $P<0.01$). Values indicated by the same letter are not statistically different, n=5. **D)** Sequence analysis of Mp*rr-myb5,2* knockout mutant lines. The wild-type *M. polymorpha* Cam-1 sequence is shown at the top, with the 20 bp gRNA target sequence highlighted with a red line. The amino acid sequence is depicted below the nucleotide sequence. **E)** Scanning electron micrograph maps of control and Mp*rr-myb2,* Mp*rr-myb5* and Mp*rr-myb5,2* thallus cross sections.

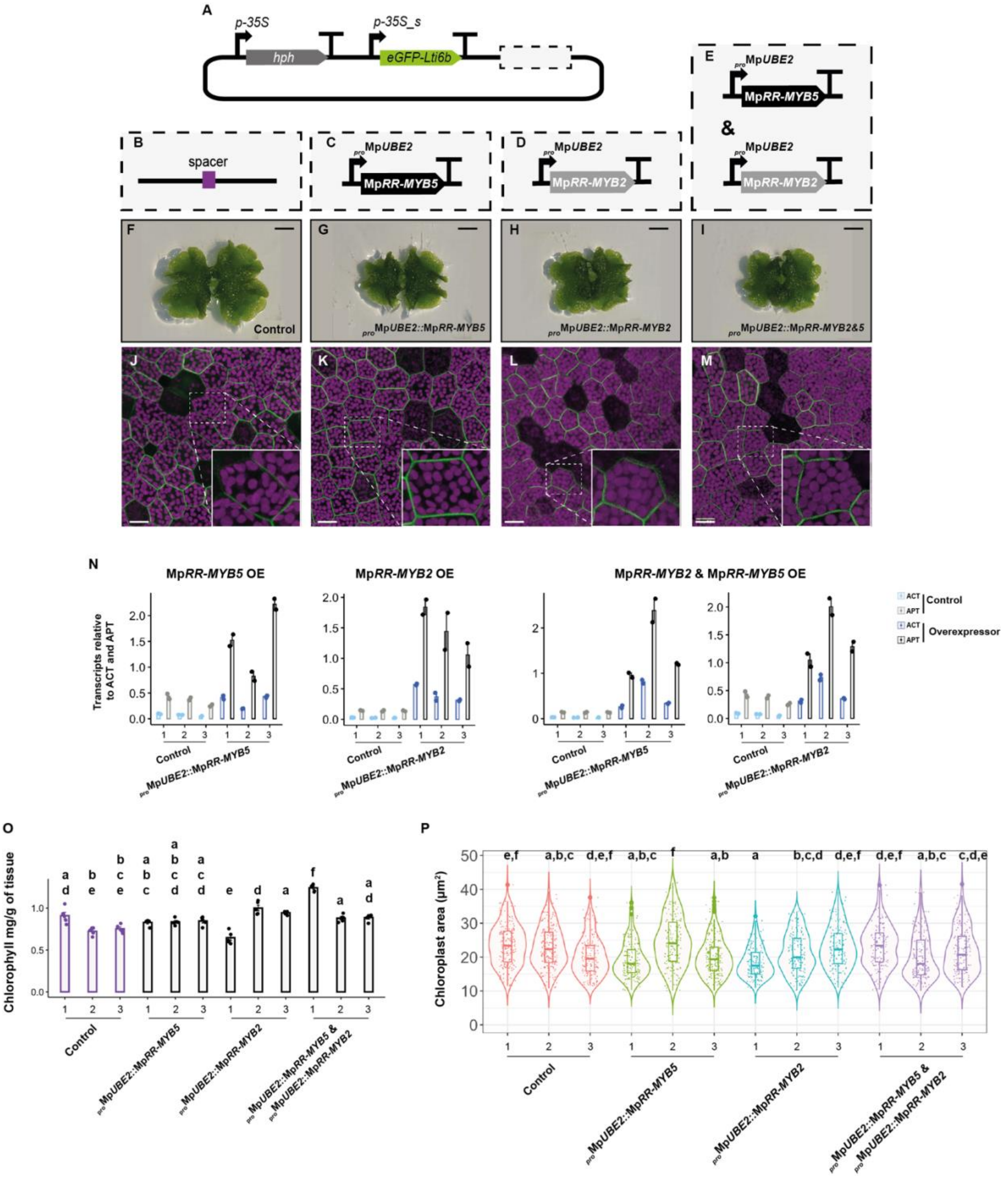
A
p-35S
hph
p-35S_s
eGFP-Lti6b
B
spacer
C
proMpUBE2
MpRR-MYB5
D
proMpUBE2
MpRR-MYB2
E
proMpUBE2
MpRR-MYB5
&
proMpUBE2
MpRR-MYB2
F
Control
G
proMpUBE2::MpRR-MYB5
H
proMpUBE2::MpRR-MYB2
I
proMpUBE2::MpRR-MYB2&5
J
K
L
M
N
MpRR-MYB5 OE
MpRR-MYB2 OE
MpRR-MYB2 & MpRR-MYB5 OE
Transcripts relative to ACT and APT
Control
proMpUBE2::MpRR-MYB5
proMpUBE2::MpRR-MYB2
ACT
APT
Control
Overexpressor
O
Chlorophyll mg/g of tissue
Control
proMpUBE2::MpRR-MYB5
proMpUBE2::MpRR-MYB2
proMpUBE2::MpRR-MYB5 &
proMpUBE2::MpRR-MYB2
P
Chloroplast area (µm²)
Control
proMpUBE2::MpRR-MYB5
proMpUBE2::MpRR-MYB2
proMpUBE2::MpRR-MYB5 &
proMpUBE2::MpRR-MYB2

**Figure S2: Mp*RR-MYB5,* Mp*RR-MYB2* and Mp*RR-MYB2 & 5* overexpression analysis**

**A-E)** Schematic representation of constructs used to overexpress Mp*RR-MYB5,* Mp*RR-MYB2* and Mp*RR-MYB2 & 5.* **F-I)** Phenotypes of Mp*RR-MYB5,* Mp*RR-MYB2* and Mp*RR-MYB2 & 5* over-expression plants. Scale bars represent 2 mm. **J-M)** Confocal microscopy images of Mp*RR-MYB5,* Mp*RR-MYB2* and Mp*RR-MYB2 & 5* over-expression plants. Scale bars represent 25 μm. **N)** qPCR analysis of Mp*RR-MYB5,* Mp*RR-MYB2* and Mp*RR-MYB2 & 5* over-expression plants. *ADENINE PHOSPHORIBOSYL TRANSFERASE 3* (*APT*) and *ACTIN 7* (*ACT*) were used as housekeeping gene controls (Saint-Marcoux et al., 2015). **O)** Barplots of chlorophyll content for the Mp*RR-MYB5,* Mp*RR-MYB2* and Mp*RR-MYB2 & 5* over-expression plants. Letters show statistical ranking using a *post hoc* Tukey test (with different letters indicating statistically significant differences at $P<0.01$). Values indicated by the same letter are not statistically different, n=5. **P)** Quantification of chloroplast area of Mp*RR-MYB5,* Mp*RR-MYB2* and Mp*RR-MYB2 & 5* over-expression plants. Letters show statistical ranking using a *post hoc* Tukey test (with different letters indicating statistically significant differences at $P<0.01$). Values indicated by the same letter are not statistically different

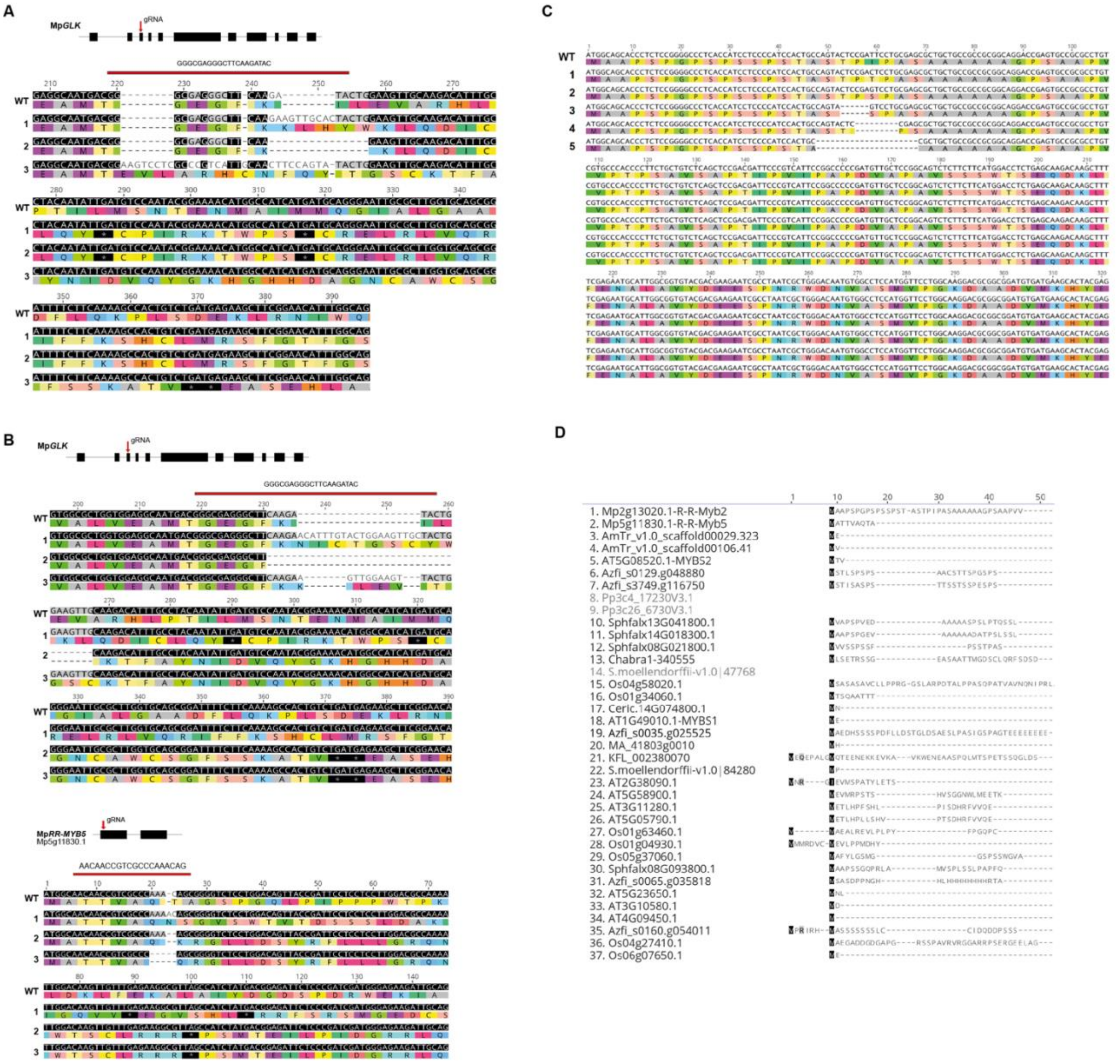
A
MpGLK
gRNA
GGGCGAGGGCTTCAAGATAC
B
MpGLK
gRNA
GGGCGAGGGCTTCAAGATAC
MpRR-MYB5
Mp5g11830.1
gRNA
AACAACCGTCGCCCAAACAG
C
D
1. Mp2g13020.1-R-R-Myb2
2. Mp5g11830.1-R-R-Myb5
3. AmTr_v1.0_scaffold00029.323
4. AmTr_v1.0_scaffold00106.41
5. AT5G08520.1-MYBS2
6. Azfi_s0129.g048880
7. Azfi_s3749.g116750
8. Pp3c4_17230V3.1
9. Pp3c26_6730V3.1
10. Sphfalx13G041800.1
11. Sphfalx14G018300.1
12. Sphfalx08G021800.1
13. Chabra1-340555
14. S.moellendorffii-v1.0|47768
15. Os04g58020.1
16. Os01g34060.1
17. Ceric.14G074800.1
18. AT1G49010.1-MYBS1
19. Azfi_s0035.g025525
20. MA_41803g0010
21. KFL_002380070
22. S.moellendorffii-v1.0|84280
23. AT2G38090.1
24. AT5G58900.1
25. AT3G11280.1
26. AT5G05790.1
27. Os01g63460.1
28. Os01g04930.1
29. Os05g37060.1
30. Sphfalx08G093800.1
31. Azfi_s0065.g035818
32. AT5G23650.1
33. AT3G10580.1
34. AT4G09450.1
35. Azfi_s0160.g054011
36. Os04g27410.1
37. Os06g07650.1

**Figure S3: CRISPR/Cas9 mediated gene editing and generation of deletions.**

**A)** Top: Schematic representation of the Mp*GLK* gene structure showing exons as black boxes. Position of gRNA is shown with an arrow. Bottom: Sequence analysis of Mp*glk* knockout mutant lines. The wild-type *M. polymorpha* Cam-1 sequence is shown at the top, with the 20 bp gRNA target sequence highlighted with a red line. The amino acid sequence is depicted below the nucleotide sequence. **B)** Top: Schematic representation of the Mp*GLK* and Mp*RR-MYB5* gene structure showing exons as black boxes. Position of gRNA is shown with an arrow. Bottom: Sequence analysis of Mp*glk,rr-myb5* double mutant lines. The wild-type *M. polymorpha* Cam-1 sequence is shown at the top, with the 20 bp gRNA target sequence highlighted with a red line. The amino acid sequence is depicted below the nucleotide sequence. **C)** Sequence analysis of Mp*glk,rr-myb5,2* knockout mutant lines. The wild-type *M. polymorpha* Cam-1 sequence is shown at the top. The amino acid sequence is depicted below the nucleotide sequence. **D)** MAFFT alignment of the N-terminus of representative RR subclass RR-MYB/CCA1-like proteins. The colouring used for that column depends on the fraction of the column that is made of letters from this group. Black: 100% similar, dark-gray: 80−100% similar, lighter gray: 60−80% similar, white: less than 60% similar.

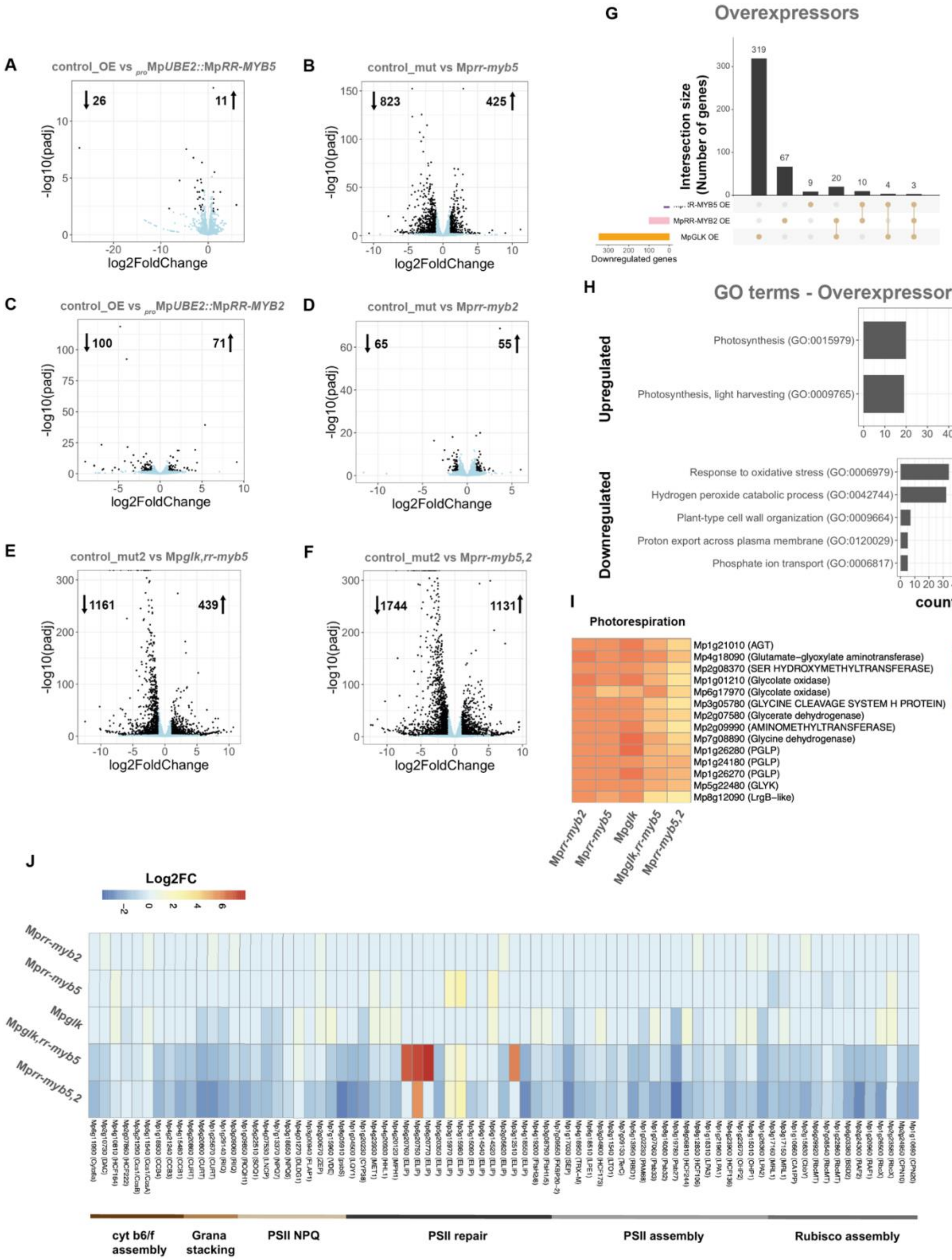
A
control_OE vs proMpUBE2::MpRR-MYB5
26
11
-log10(padj)
log2FoldChange
B
control_mut vs Mprr-myb5
823
425
C
control_OE vs proMpUBE2::MpRR-MYB2
100
71
D
control_mut vs Mprr-myb2
65
55
E
control_mut2 vs Mpglk,rr-myb5
1161
439
F
control_mut2 vs Mprr-myb5,2
1744
1131
G
Overexpressors
Intersection size (Number of genes)
319
67
9
20
10
4
3
MpRR-MYB2 OE
MpGLK OE
Downregulated genes
H
GO terms - Overexpressors
Upregulated
Photosynthesis (GO:0015979)
Photosynthesis, light harvesting (GO:0009765)
Downregulated
Response to oxidative stress (GO:0006979)
Hydrogen peroxide catabolic process (GO:0042744)
Plant-type cell wall organization (GO:0009664)
Proton export across plasma membrane (GO:0120029)
Phosphate ion transport (GO:0006817)
counts
I
Photorespiration
Mp1g21010 (AGT)
Mp4g18090 (Glutamate−glyoxylate aminotransferase)
Mp2g08370 (SER HYDROXYMETHYLTRANSFERASE)
Mp1g01210 (Glycolate oxidase)
Mp6g17970 (Glycolate oxidase)
Mp3g05780 (GLYCINE CLEAVAGE SYSTEM H PROTEIN)
Mp2g07580 (Glycerate dehydrogenase)
Mp2g09990 (AMINOMETHYLTRANSFERASE)
Mp7g08890 (Glycine dehydrogenase)
Mp1g26280 (PGLP)
Mp1g24180 (PGLP)
Mp1g26270 (PGLP)
Mp5g22480 (GLYK)
Mp8g12090 (LrgB−like)
Mprr-myb2
Mprr-myb5
Mpglk
Mpglk,rr-myb5
Mprr-myb5,2
J
Log2FC
Mprr-myb2
Mprr-myb5
Mpglk
Mpglk,rr-myb5
Mprr-myb5,2
cyt b6/f assembly
Grana stacking
PSII NPQ
PSII repair
PSII assembly
Rubisco assembly

**Figure S4: *M. polymorpha* RNA sequencing analysis.**

**A-B)** Volcano plots showing differentially expressed genes in Mp*RR-MYB5* overexpression vs control plants and for the Mp*rr-myb5* mutant plants. Blue dots indicate genes with a padj-value ≥ 0.01, while the black ones have a padj-value ≤ 0.01. The total number of differentially expressed genes (DEGs) are indicated at the top of the graph. **C-D)** Volcano plots showing differentially expressed genes Mp*RR-MYB2* overexpression vs control plants and for the Mp*rr-myb2* mutants plants vs control. Blue dots indicate genes with a padj-value ≥ 0.01, while the black ones have a padj-value ≤ 0.01. The total number of DEGs are indicated at the top of the graph. **E)** Volcano plots showing differentially expressed genes for the Mp*glk,rr-myb5* mutants plants vs control. **F)** Volcano plots showing differentially expressed genes for the Mp*rr-myb5,2* mutants plants vs control. **G)** Upset diagrams showing shared downregulated genes in the Mp*RR-MYB5,* Mp*RR-MYB2* and Mp*GLK* over-expression plants **H)** Gene ontology term enrichment Mp*GLK* over-expression plants. **I)** Heatmap of differentially expressed photorespiration genes in Mp*rr-myb2,* Mp*rr-myb5,* Mp*glk* as well as Mp*glk,rr-myb5* and Mp*rr-myb5*,*2* double mutants. **J)** Heatmap of differentially expressed photosynthesis associated genes in Mp*rr-myb2,* Mp*rr-myb5,* Mp*glk* as well as Mp*glk,rr-myb5* and Mp*rr-myb5*,*2* double mutants.

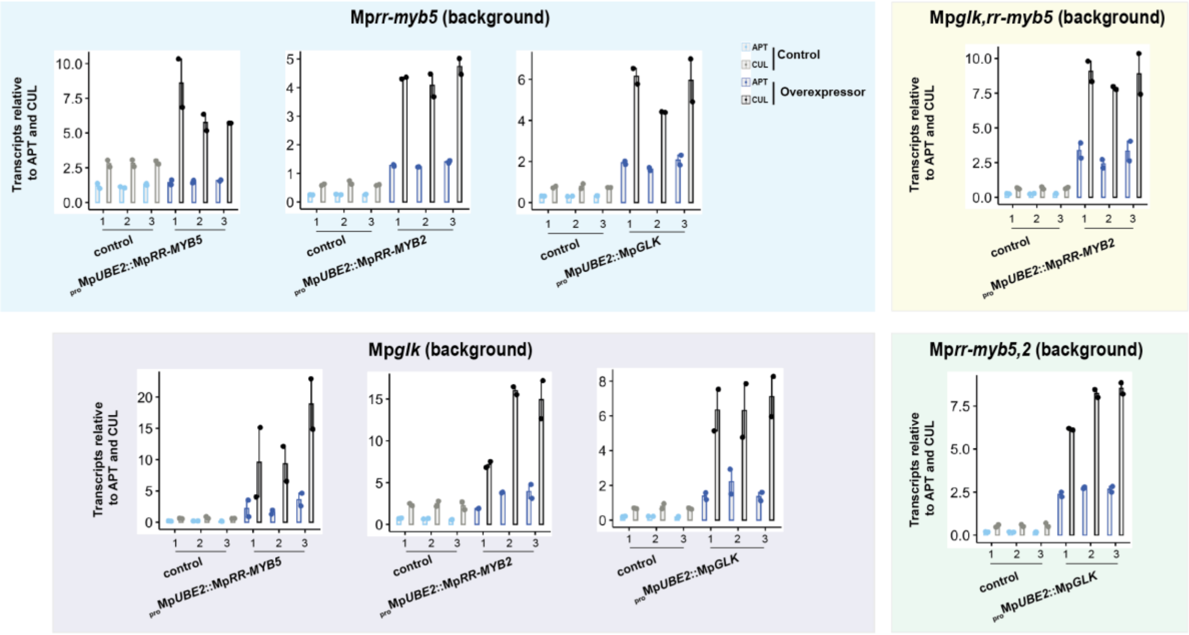


**Figure S5: Quantitative polymerase chain reactions for transgene over-expression confirmation.**

qPCR analysis of Mp*rr-myb5,* Mp*glk,* Mp*glk,rr-myb5* and Mp*rr-myb5,2* mutants complemented with Mp*RR-MYB5,* Mp*RR-MYB2* and Mp*GLK* driven by the Mp*UBE2* promoter*. ADENINE PHOSPHORIBOSYL TRANSFERASE 3* (*APT*) and *CULLIN 1* (*CUL*) were used as housekeeping gene controls (Saint-Marcoux et al., 2015).

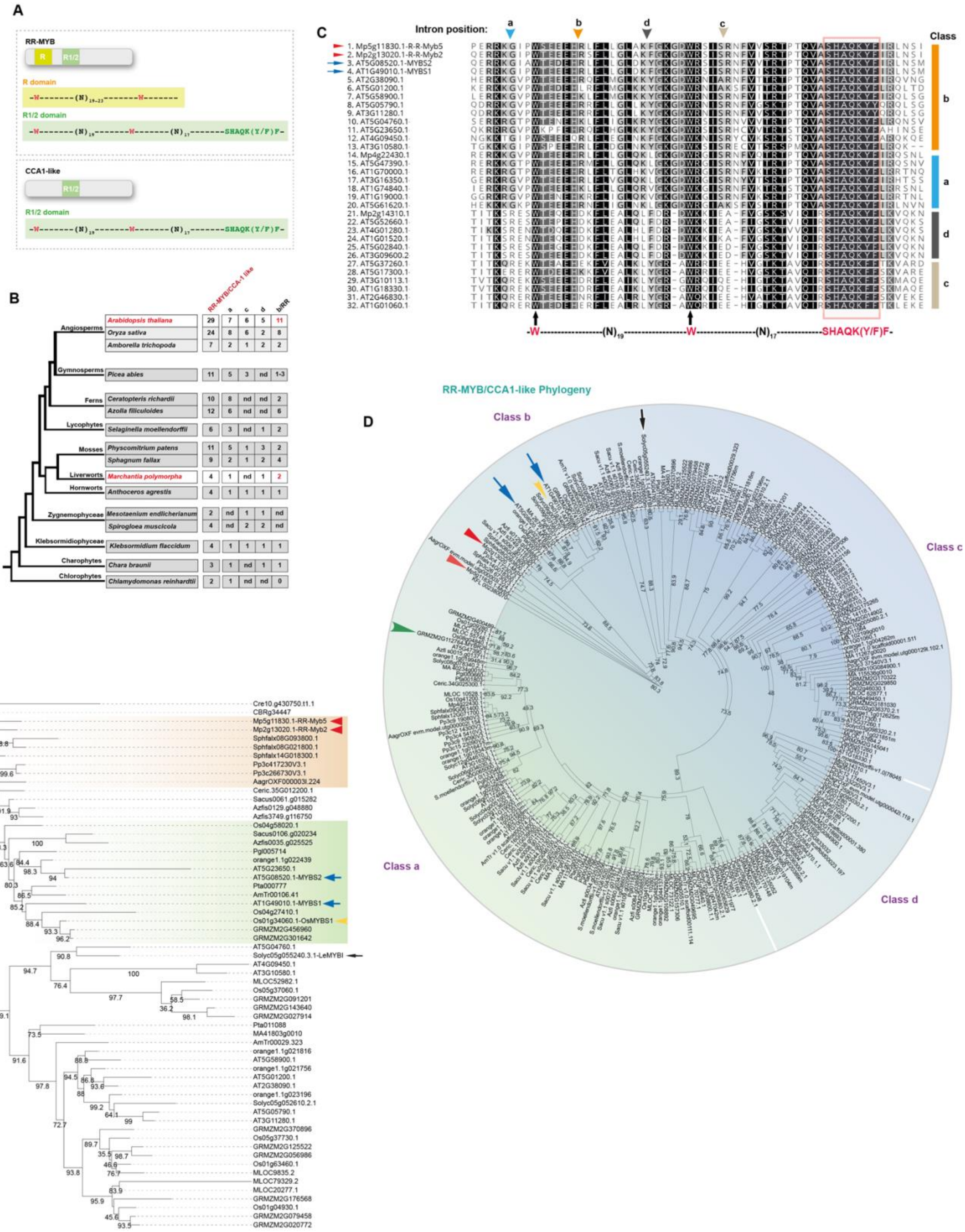
A
RR-MYB
R domain
R1/2 domain
CCA1-like
R1/2 domain
B
Angiosperms
Arabidopsis thaliana
Oryza sativa
Amborella trichopoda
Gymnosperms
Picea abies
Ferns
Ceratopteris richardii
Azolla filiculoides
Lycophytes
Selaginella moellendorffii
Mosses
Physcomitrium patens
Sphagnum fallax
Liverworts
Marchantia polymorpha
Hornworts
Anthoceros agrestis
Zygnemophyceae
Mesotaenium endlicherianum
Spirogloea muscicola
Klebsormidiophyceae
Klebsormidium flaccidum
Charophytes
Chara braunii
Chlorophytes
Chlamydomonas reinhardtii
C
Intron position:
Class
D
RR-MYB/CCA1-like Phylogeny
Class b
Class c
Class a
Class d
E

**Figure S6: The RR-MYB/CCA1-like related transcription factor family.**

**A)** Schematic representation of RR-MYB/CCA1-like protein structure. Mp*RR-Myb5* belongs to the RR-MYB/ CIRCADIAN CLOCK ASSOCIATED1 - like (CCA1-like) subfamily of the MYB-related transcription factors. The RR-MYB/CCA1-like subfamily contains two different classes of genes, the RR and the CCA1-like. They both contain a Myb domain with a characteristic SHAQK(Y/F)F DNA binding motif called R1/2, but the RR class has an additional MYB-like domain called R. Conserved "W" residues and SHAQK(Y/F)F domain are indicated at the bottom of the alignment. The CCA1-like class can further be categorised into three additional groups based on the intron position within the R1/2 domain, called a, c and d (Du et al. 2013). Intron position within the R1/2 domain of the RR class is also distinct from the three intron positions found in the CCA1-like class (b group) and can serve as an additional character/evidence for RR ortholog identification. *A. thaliana* has a total of 29 RR-MYB/CCA1-like genes, of which seven belong to the a class, six to the c class, five to the d class and 11 to the b class/RR. Rice has a total of 25 RR-MYB/CCA1-like genes, of which eight belong to the a class, six to the c class, nine to the d class and nine to the b class/RR. *M. polymorpha* has a total of only four RR-MYB/CCA1-like genes, of which one belongs to the a class, one to the d class and two to the b class/RR. The c class seems to be absent. **B)** Plant phylogeny with representative species for which whole genome assemblies are available. RR-MYB/CCA1-like gene numbers are shown (nd: not detected). **C)** MAFFT alignment of R1/2 domain of *M. polymorpha* and *A. thaliana* RR-MYB/CCA1-like transcription factor. *M. polymorpha and A. thaliana* proteins examined in this study are indicated with red arrowheads and blue arrows respectively. Positions of introns are marked above the alignment with arrowheads. The colouring used for that column depends on the fraction of the column that is made of letters from this group. Black: 100% similar, dark-gray: 80−100% similar, lighter gray: 60−80% similar, white: less than 60% similar. **D)** RR-MYB/CCA1-like related phylogeny. Numbers on branches represent SH-aLRT test support (Guindon et al. 2010). *M. polymorpha*, *A. thaliana*, Rice and Tomato proteins are indicated with red arrowheads, blue arrows, yellow arrowhead and black arrow respectively. **E)** RR-MYB related phylogeny. Numbers on branches represent ultrafast bootstrap support (Hoang et al. 2018). *M. polymorpha*, *A. thaliana*, Rice and Tomato proteins are indicated with red arrowheads, blue arrows, yellow arrowhead and black arrow respectively.

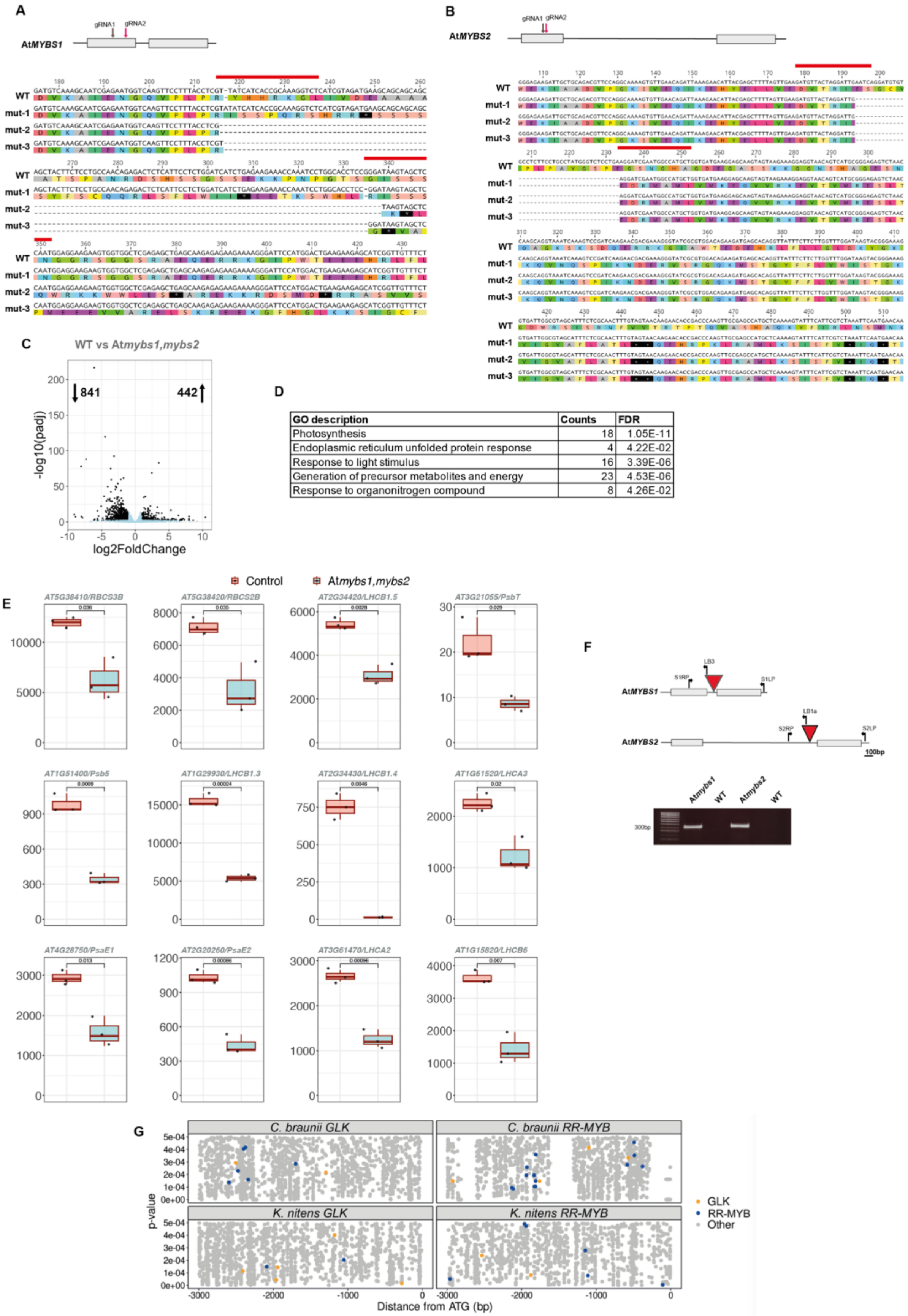


| GO description | Counts | FDR |
|---|---|---|
| Photosynthesis | 18 | 1.05E-11 |
| Endoplasmic reticulum unfolded protein response | 4 | 4.22E-02 |
| Response to light stimulus | 16 | 3.39E-06 |
| Generation of precursor metabolites and energy | 23 | 4.53E-06 |
| Response to organonitrogen compound | 8 | 4.26E-02 |

**Figure S7: CRISPR/Cas9 mediated gene editing and generation of deletions.**

**A-B)** Top: Schematic representation of the At*MYBS1* and At*MYBS2* gene structure showing exons as grey boxes. Positions of gRNAs are shown with a grey/red arrow. Bottom: Sequence analysis of At*mybs1* and At*mybs2* knockout double mutant lines. The wild-type *A. thaliana Col-0* sequence is shown at the top, with the 20 bp gRNA target sequence highlighted with a red line. The amino acid sequence is depicted below the nucleotide sequence. **C)** Volcano plot showing differentially expressed genes in At*mybs1,mybs2* mutants. Blue dots indicate genes with a padj-value ≥ 0.01, while the black ones have a padj-value ≤ 0.01. The total number of DEGs are indicated at the top of the graph. **D)** Top five GO terms estimated using PANTHER (GO-Slim Biological Processes). **E)** Boxplots for representative downregulated photosynthesis associated genes in At*mybs1,mybs2* mutants*.* Data are presented as Transcripts per Million (TPM) values. P-values of two-tailed *t*-test are shown. **F)** Genotyping of At*mybs1* (SAIL_1184_D04) and At*mybs2* (SALK_150774) mutant lines. Top: Schematic representation of At*MYBS1* and At*MYBS2* gene structure showing exons as grey rectangles. The T-DNA insertion positions are indicated with red arrowheads (not in scale). The positions of the primer sequences used for genotyping are shown with bended arrows. Primer pairs S1RP & LB3 and S2RP & LB1a were used to confirm the presence of the T-DNA insert in At*mybs1* and At*mybs2* mutants respectively. Bottom: PCR analysis for the confirmation of T-DNA insert presence in mutants. **G)** Scatter plots showing position and predicted binding affinity of GLK, RR-MYB binding and other motifs 3000 bp upstream of the translation start site of *Chara braunii* and *Klebsormidium flaccidum* GLK and RR-MYB putative homologs. Y-axis shows p-values of matches between upstream regions and motif position weight matrices, and x-axis shows position of the motif center relative to the transcriptional start site.

**ImageJ macro:**

```
run("Duplicate...");
run("Smooth");
run("Auto Local Threshold", "method=Phansalkar radius=5 parameter_1=0 parameter_2=0 white");
run("Watershed");
run("Analyze Particles...", "size=1-500 circularity=0.01-1.00 display exclude clear add");
```
